# Disruption of the ganglioside-plasma membrane calcium ATPase-neuroplastin nexus leads to impaired calcium homeostasis in glioblastoma

**DOI:** 10.64898/2025.12.12.693888

**Authors:** Borna Puljko, Anja Kafka, Nikolina Maček Hrvat, Anja Bukovac, Niko Njirić, Antonia Jakovčević, Vinka Potočki, Fran Dumančić, Ana Karla Vodanović, Ana Ujevic, Eva Josic, Svjetlana Kalanj-Bognar, Kristina Mlinac-Jerkovic, Nives Pećina-Šlaus

**Affiliations:** Laboratory for Molecular Neurobiology and Neurochemistry, Croatian Institute for Brain Research, School of Medicine, University of Zagreb, Zagreb, Croatia; Department of Chemistry and Biochemistry, School of Medicine, University of Zagreb, Zagreb, Croatia; Laboratory for Neurooncology, Croatian Institute for Brain Research, School of Medicine, University of Zagreb, Zagreb, Croatia; Department of Biology, School of Medicine, University of Zagreb, Zagreb, Croatia; Institute for Medical Research and Occupational Health, Zagreb, Croatia; Department of Neurosurgery, University Hospital Center Zagreb, Zagreb, Croatia; Department of Pathology and Cytology Ljudevit Jurak, University Hospital Center Sestre Milosrdnice, Zagreb, Croatia; Department of Clinical Neuroscience, Karolinska Institutet, Stockholm, Sweden

## Abstract

Glioblastoma (GBM), an exceedingly invasive brain tumor, is characterized by the disruption of multiple signaling pathways, particularly those governing calcium homeostasis. Additionally, cell membranes in GBM exhibit altered lipidomic profiles, especially complex sialoglycans gangliosides, which affect plasma membranes both structurally and functionally. In this work we show a disruption of a plasma membrane molecular triad composed of gangliosides, neuroplastin (Np), and plasma membrane calcium ATPase (PMCA) in GBM, leading to a severely diminished PMCA activity. Our transcriptomic, epigenetic, proteomic, lipidomic, and *in silico* analyses consistently demonstrated downregulation of Np and PMCA isoforms both at mRNA and protein levels in our GBM cohort, in agreement with a broader trend observed in public datasets. Analysis of the uniquely changed GBM ganglioside profile in the context of PMCA and Np expression and PMCA activity reveals GD3 as a key molecular determinant of reduced PMCA activity in GBM samples. In contrast, GD1b, necessary for optimal PMCA function in healthy tissue, is significantly reduced in GBM. Our findings establish the gangliosides-PMCA-Np nexus as a crucial complex involved in calcium signaling in GBM, contributing to GBM etiology and progression.

## Introduction

Glioblastoma multiforme (GBM) is the most common and most aggressive grade 4 astrocytoma in adults, with an overall survival of approximately 15 months and a recurrence-free survival below <7% at 5 years, despite aggressive multimodal therapy ^1,2^. GBM is characterized by rapid growth, diffuse infiltration of the brain parenchyma, local resistance to treatment, and a significant capacity to alter its microenvironment ^3^. A central hallmark of GBM progression is the ability to hijack cell signaling pathways, promoting tumor growth, stemness, and immune evasion ^4–6^. Although several different molecular pathways are altered in GBM, disruption of calcium homeostasis has emerged as one of the central drivers of malignancy, as it is responsible for increased motility, survival in a hypoxic environment, and resistance to apoptosis, highlighting it as an essential target for investigation ^7–10^.

The lipid composition of cell membranes plays a crucial role in regulating cellular physiology, particularly in signaling, adhesion, and migration, which are pivotal in determining tumor etiology and behavior ^11,12^. Alteration in lipid composition is also a hallmark of cancer cells, and GBM is no exception. Elevated concentrations of cholesterol, sphingolipids, and specific ganglioside species are reported in GBM cell membranes ^13,14^. These variations in lipid content and composition modify membrane fluidity, curvature, and structure, influencing the localization and function of membrane-associated proteins, including calcium-handling proteins such as plasma membrane calcium ATPase (PMCA) ^15–19^.

Gangliosides, a class of silalylated glycosphingolipids primarily concentrated in the central nervous system, exhibit a changed structural and compositional pattern in GBM, with increased levels of GD3 and GD2 being hallmark features ^20–23^. These gangliosides are associated with enhanced primary tumor growth, migration, immune evasion, and resistance to apoptosis ^23,24^. Additionally, gangliosides affect the function of critical calcium pumps, such as PMCA, thereby potentially compromising calcium homeostasis in glioblastoma cells by altering their stability, localization, and functionality ^25^. However, how these lipid perturbations translate into calcium dysregulation and promote GBM aggressiveness remains poorly understood.

Neuroplastin (Np) is a transmembrane protein belonging to the immunoglobulin superfamily, important for synaptic plasticity, neurodevelopment, and neurodegeneration^26–32^; however, its role in gliomagenesis has not yet been described. Np serves as an auxiliary subunit of PMCAs and is crucial for their correct functionality and trafficking to the plasma membrane ^33,34^. The Np-PMCA complex is an essential regulator of intracellular calcium homeostasis ^35,36^. PMCA facilitates the extrusion of calcium from the cytoplasm maintaining physiological levels vital for cellular survival and signaling. Without Np, PMCAs are unable to perform their role in calcium handling efficiently ^33,34,37^. While differences in Np expression have been identified in various cancers including lung, breast, and colorectal cancers ^38–47^, it has not been a focus of study in GBM. Both Np and PMCA exist in several isoforms. Np, coded by the *NPTN* gene, has two isoforms, Np55 and Np65, arising from alternative splicing ^30^. PMCA has four main isoforms (PMCA1, PMCA2, PMCA3 and PMCA4) coded by the separate genes *ATP2B1*, *ATP2B2*, *ATP2B3* and *ATP2B4*, respectively. These four main isoforms generate over 30 variants through alternative splicing, adding an additional level of complexity to their investigation ^48^. *ATP2B3* (PMCA3) was excluded from our analyses due to its localization on the X chromosome in humans^49^.

Here, we show a disruption of a plasma membrane molecular nexus composed of gangliosides, Np, and PMCA in GBM, resulting in severely diminished PMCA activity, likely contributing to GBM etiology. Our multi-level experimental and *in silico* approach examined the protein and mRNA expression levels of PMCA and Np isoforms, their epigenetic profiles, PMCA activity and relevant ganglioside expression patterns to delineate how this molecular triad could contribute to calcium extrusion impairment in GBM. Gaining insight into the disturbance of normal associations between gangliosides, PMCA and Np in GBM may lay the groundwork for future treatment approaches targeting this molecular network.

## Results

### Coordinated downregulation and mutational landscape of neuroplastin and PMCA isoforms in glioblastoma

To assess the gene expression of PMCA isoforms and Np isoforms, we performed quantitative real-time polymerase chain reaction (qRT-PCR) on resected GBM tissue and age-matched healthy cerebrocortical samples (Fig. 1.a).

**Fig. 1.**
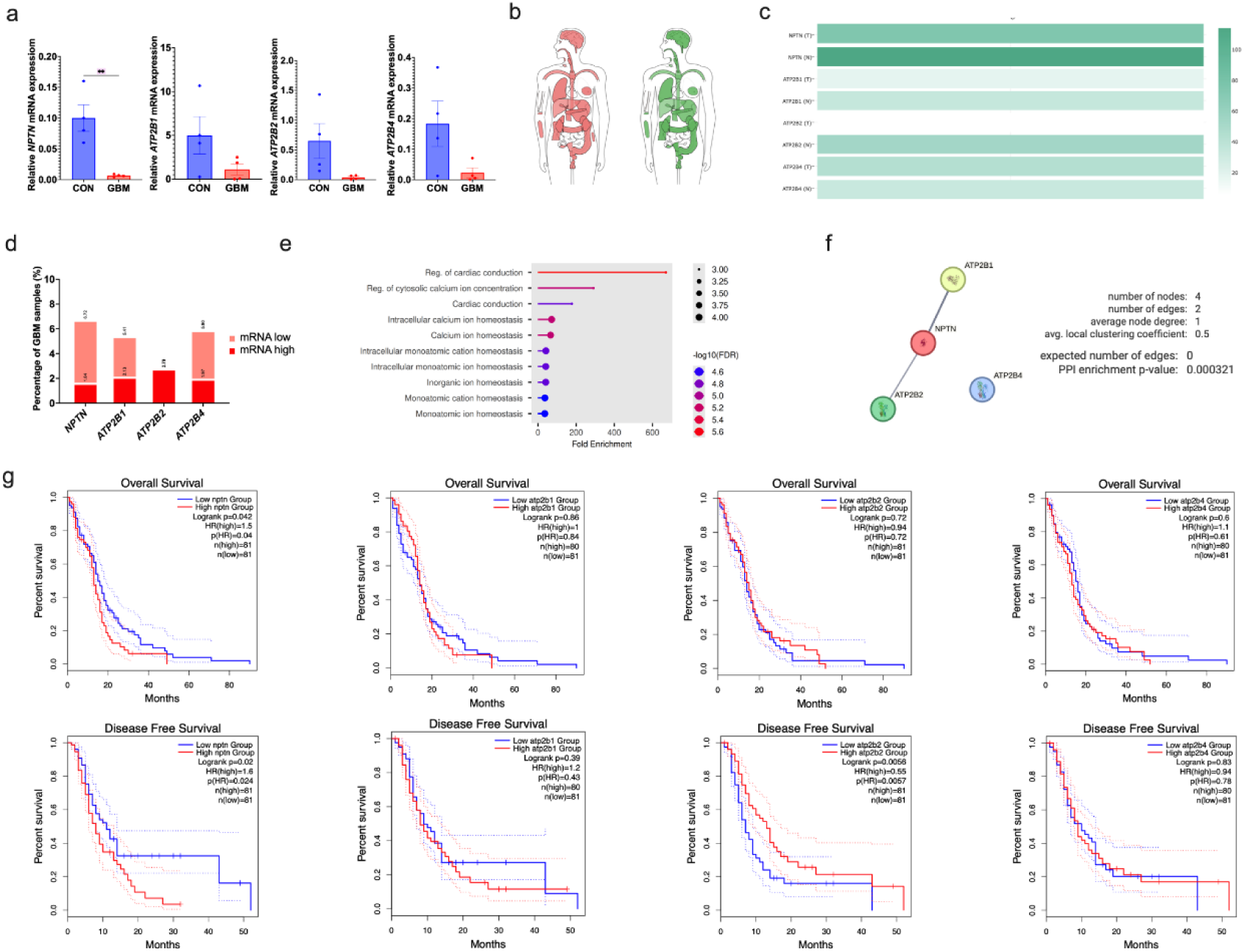
mRNA expression analysis of Np and PMCA in glioblastoma and the progonostic significance of the altered gene expression patterns. a Quantitative PCR analysis of resected glioblastoma (GBM) and non-tumor brain samples (CON; controls). Data is shown as fold change mean ± SEM. ** = p < 0,05; two-tailed unpaired *t*-test; n = 4 per group. b Tissue distribution of Np coding gene (*NPTN)* expression in tumors (left, red) and healthy tissues (right, green) visualized using GEPIA BodyMap. The median expression of the *NPTN* gene is presented on a log₂(TPM+1) scale. c *In silico* comparison of mRNA expression levels from TCGA datasets of *NPTN* and PMCA isoforms genes in GBM tumor (n = 163) versus normal brain tissue (n = 207) from GEPIA2. Bar plot shows expression levels for *NPTN*, *ATP2B1*, *ATP2B2*, and *ATP2B4* in tumor (T) and healthy normal (N) samples from the TCGA/GTEx matched dataset. d Distribution of GBM samples (n = 163) by mRNA expression levels of *NPTN*, *ATP2B1*, *ATP2B2*, *ATP2B* genes from the TCGA GBM cohort (Firehose Legacy). The bar chart shows the proportion of tumor samples classified as “low” (below z-score −2) and “high” (above z-score +2) expressing. cBioPortal was used to stratify samples based on mRNA expression levels. e GO Biological Process enrichment analysis of the Np-PMCA gene cluster using ShinyGO. Enriched pathways (FDR < 0.05) are visualized as lollipop plots, ranked by fold enrichment. Circle size represents gene contributions, and color denotes significance (red: lower FDR). f STRING protein-protein interaction network between *NPTN* and PMCA isoforms genes (*ATP2B1*, *ATP2B2*, *ATP2B4*). Edges represent evidence-based associations, such as co-expression, experimental data, and curated databases. g Kaplan-Meier survival curves from GEPIA showing overall and disease-free survival in GBM patients stratified by *NPTN*, *ATP2B1*, *ATP2B2*, and *ATP2B4* expression. Log-ranked p values, HR, and group sizes are provided.

*NPTN* mRNA levels (encompassing both Np isoforms, Np55 and Np65) were markedly lower in GBM relative to control cerebrocortical samples (p = 0.0042, t = 3.523, dF = 12, two-tailed unpaired t-test), signifying significant downregulation. The expression of *ATP2B1*, *ATP2B2*, and *ATP2B4* coding for PMCA isoforms, was also consistently decreased in GBM relative to controls; however, statistical significance was not attained, likely due to the limited sample size. To further validate these results, we employed GEPIA2, an interactive tool that amalgamates extensive TCGA and GTEx datasets. Fig. 1.b shows the median mRNA expression of *NPTN* in tumor and paired normal tissues from BodyMap, indicating which tumors have so far been identified as having altered *NPTN* expression. Fig. 1.c. reveals an *in silico* comparison of mRNA expression levels for *NPTN* and PMCA isoforms genes in GBM compared to normal brain tissue. This analysis encompassing a larger sample size clearly shows that the Np expression differences are extensively present in various tumor types, accompanied by markedly different mRNA levels of PMCA isoforms genes, especially *ATP2B2*, which is all in agreement with our qRT-PCR results. *ATP2B4* was almost unchanged between tumorous and non-tumorous large-scale RNAseq sets. Furthermore, we examined the percentage of GBM samples with low expression levels using cBioPortal TCGA dataset (Fig. 1d). *NPTN*, *ATP2B1*, and *ATP2B4* were downregulated more frequently, whereas *ATP2B2* showed a higher expression profile in cohort of GBM tumors.

To better understand the functional significance of the Np-PMCA network, we performed Gene Ontology (GO) enrichment analysis using the ShinyGO 0.85 platform. Analysis of the four-gene set revealed strong enrichment for biological processes related to ion homeostasis and calcium signaling (Fig. 1.e). To explore potential physical and functional relationships between Np and PMCA isoforms, we analyzed protein-protein interactions (PPI) using the STRING analysis tool (Fig. 1.f). Np interacted with two plasma membrane Ca²⁺ ATPase isoforms, *ATP2B1* (PMCA1) and *ATP2B2* (PMCA2), supporting a high-confidence functional interaction network. The PPI network exhibited substantial enrichment (p = 0.000321), indicating non-random biological associations among these Ca^2+^-handling proteins. However, *ATP2B4* (PMCA4) did not interact with Np. This is in line with data reporting that Np controls certain PMCA isoforms to govern calcium efflux.

Next, we investigated the potential prognostic significance of the notable co-expression patterns between *NPTN* and genes for PMCA isoforms using Kaplan-Meier survival analysis with GEPIA2 (Fig.1.g). Kaplan-Meier curves demonstrated that higher *NPTN* expression was associated with lower overall survival (HR = 1.5, p = 0.04) and disease-free survival (HR = 1.6, p = 0.024). Notably, high A*TP2B2* expression among PMCA isoforms was significantly associated with favorable disease-free survival (HR = 0.55, p = 0.0057); no significant associations were observed for *ATP2B1* or *ATP2B4*. To further evaluate genomic alterations affecting the PMCA and Np, we thoroughly analyzed somatic mutations in *NPTN*, *ATP2B1*, *ATP2B2*, and *ATP2B4* using the cBioPortal and COSMIC databases (Supplementary Tables 1-4). Across 4,466 samples in total from 11 glioma and glioblastoma studies (Pediatric Brain Cancer (218), Diffuse Glioma (693), Diffuse Glioma (444), Glioma (1004), Glioma (91), IDH-mutated Diffuse Glioma (73), Low-Grade Gliomas (61), Merged Cohort of LGG and GBM (1122), Glioblastoma (99), Glioblastoma (42), and Glioblastoma Multiforme (619), COSMIC reported 12 *NPTN* gene mutations, all missense (Supplementary Table 1). The *ATP2B1* gene harbored mutations in 22 samples: 18 missense, 1 deletion, 1 silent, and 2 substitutions with unknown consequences (Supplementary Table 2). The *ATP2B2* gene showed the highest mutation rate, with 39 glioma samples—27 missense and 12 same-sense substitutions (Supplementary Table 3). The *ATP2B4* gene harboured mutations in 27 samples: 17 missense, 2 intronic, 2 nonsense, and 6 silent substitutions (Supplementary Table 4). *In silico* predictions using MutationTaster, PolyPhen-2, and SIFT on canonical sequences classified multiple variants as probably damaging or disease-causing, notably *NPTN* p.L231Q, p.R156H, p.E99K, p.T78N; *ATP2B1* p.N1005S, p.K883N, p.W844R, p.A817D, p.A804V, p.D631V, p.D174G, p.L168I, p.S141F; *ATP2B2* p.K1163R, p.G679S, p.D454N, p.R437C, p.G413E, p.V390I, p.V390I, p.D143, p.A114V, p.D454N and p.A914G, and for *ATP2B4* p.P139T, p.R178P, p.S249F, p.L380I, p.L503F, p.K561N, p.R587H, suggesting functional impairment. Full details of mutation type, frequency, and predicted pathogenicity are provided in Supplementary Tables 1-4. The consistent lower expression of *NPTN* and specific PMCA genes, potentially caused by damaging mutations, suggests a convergent disruption of calcium extrusion mechanisms in glioblastoma.

### Correlation between gene expression of Np, PMCA isoforms and ganglioside biosynthesis genes

To explore transcriptional correlations between *NPTN*, PMCA isoforms genes, and key ganglioside biosynthetic pathway genes (Fig. 2.a), we performed Pearson correlation co-expression analysis amongst 207 TCGA GBM sample datasets using GEPIA2 (Fig. 2.b to 2.f).

**Fig. 2.**
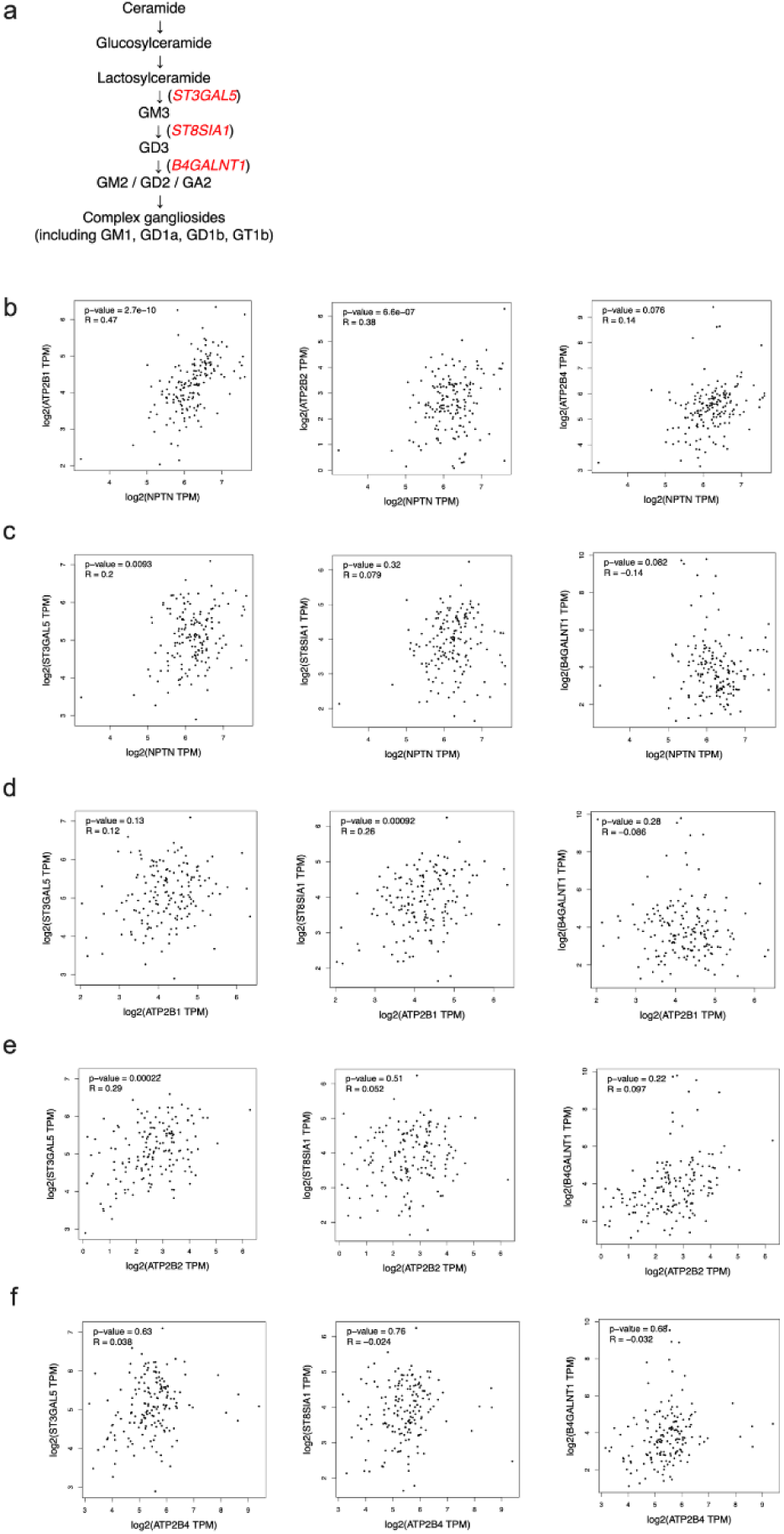
Co-expression analysis of PMCA isoforms, Np isoforms and ganglioside biosynthetic enzyme coding genes. **a** A simplified schematic representation of the ganglioside biosynthesis pathway. Enzymes of interest in this study are marked in red. **b-e** Co-expression analysis of *NPTN*, PMCA isoforms genes, and ganglioside biosynthetic enzyme genes in GBM. Scatter plots from GEPIA2 show Pearson correlation of gene expression between selected genes in the GBM sample (n = 163) on a log₂(TPM+1) scale. R and p values are indicated. Correlation is shown for: **b** *NPTN* and PMCA isoform genes *ATP2B1*, *ATP2B2* and *ATP2B4*; **c** *NPTN* and genes coding for ganglioside biosynthesis enzymes (*ST3GAL5, St8SIA1, B4GALNT1*); **d** PMCA1 gene and genes coding for ganglioside biosynthesis enzymes; **e** PMCA2 gene and genes coding for ganglioside biosynthesis enzymes; **f** PMCA4 gene and genes coding for ganglioside biosynthesis enzymes.

*NPTN* expression showed positive correlations with *ATP2B1* (R = 0.47, p = 2.7 × 10⁻¹⁰) and *ATP2B2* (R = 0.38, p = 6.6 × 10⁻⁷). The correlation with *ATP2B4* was weak and not significant (R = 0.14, p = 0.076).

Among the genes that code for ganglioside biosynthetic enzymes, *ST3GAL5* (which codes for GM3 synthase) had a weak positive correlation (R = 0.20, p = 0.0093). In contrast, *ST8SIA1* (coding for GD3 synthase) and *B4GALNT1* (coding for GM2/GD2 synthase) did not have a significant correlation with *NPTN* (r = 0.079, p = 0.32 and r = −0.14, p = 0.082, respectively). Among PMCA isoforms-coding-genes, *ATP2B1* exhibited a notably weak positive connection with *ST8SIA1* (r = 0.26, p = 0.00092), while showing no significant correlation with *ST3GAL5* (R = 0.12, p = 0.13) and *B4GALNT1* (r = −0.086, p = 0.28). *ATP2B2* had a positive connection with *ST3GAL5* (r = 0.29, p = 0.00022), but no significant correlations with *ST8SIA1* (r = 0.052, p = 0.51) or *B4GALNT1* (r = 0.097, p = 0.22). There was no significant association between *ATP2B4* and any of the enzymes of ganglioside biosyntheis.

### Epigenetic modulation of *NPTN* and PMCA genes expression in glioblastoma

Given the coordinated downregulation of *NPTN* and PMCA isoform genes observed in our GBM samples and in public databases, we examined the upstream regulatory mechanisms governing this transcriptional regulation. DNA methylation and transcription factor (TF) activity are essential for regulating gene expression in cancer. We utilized both experimental and computational approaches to evaluate the epigenetic and transcriptional landscape of *NPTN* and PMCA genes (Fig. 3).

**Fig. 3.**
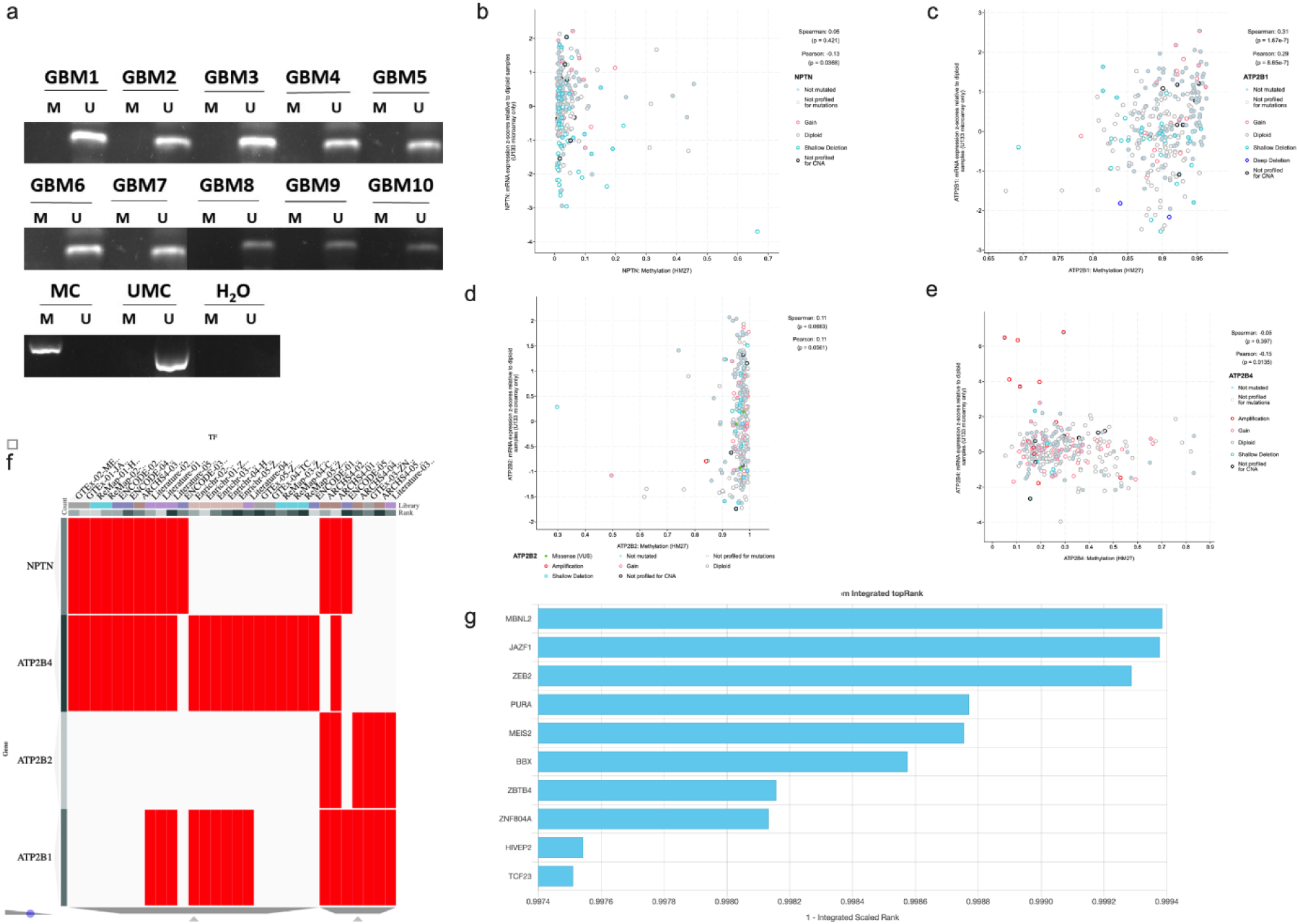
Analysis of promoter methylation, correlation to mRNA expression and transcription factor (TF) and gene interactions analysis for *NPTN* and PMCA isoforms genes. a Methylation-specific PCR (MSP) analysis of the *NPTN* promoter region in 10 glioblastoma samples. Presence of PCR bands in the U lane indicates unmethylated alleles; M: methylated alleles. MC: methylated control; UMC: unmethylated control; H₂O: negative control. **b-e** Correlation between promoter methylation and mRNA expression in GBM samples (TCGA, cBioPortal) for: b *NPTN*; c *ATP2B1*; d *ATP2B2*, and e *ATP2B4*.

Pearson and Spearman coefficients are shown, n = 278 GBM samples. **f** A clusterogram of transcription factor (TF) and gene interactions across ChEA3 datasets for *NPTN* and PMCA isoforms genes. Red bars denote TF and gene associations based on ChIP-seq, GTEx, ENCODE, ReMap, and literature. **g** Top 10 transcription factors ranked by integrated scaled score (ChEA3 TopRank), showing candidate TFs co-regulating *NPTN* and PMCA genes.

The results of the methylation-specific PCR (MSP) we performed for *NPTN* promoter regions showed clear unmethylated PCR products in all ten GBM samples (Fig. 3.a), indicating that promoter hypermethylation is not the primary mechanism of *NPTN* downregulation in this cohort. Methylation events of the queried genes were also investigated *in silico*. Correlation analysis between DNA methylation and mRNA expression in TCGA GBM samples (n = 278) revealed distinct methylation–expression relationships across the queried genes. For *NPTN* (Fig. 3.b), no significant correlation was observed (Spearman’s ρ = 0.05, p = 0.421; Pearson’s r = –0.13, p = 0.0368) suggesting that its expression is largely methylation-independent. In contrast, *ATP2B1* displayed a moderate and statistically significant positive correlation between methylation and mRNA expression (Spearman’s ρ = 0.31, p = 1.6 × 10⁻⁷; Pearson’s r = 0.29, p = 8.6 × 10⁻⁷) (Fig. 3.c). For *ATP2B2*, a non-significant positive correlation was detected (Spearman’s ρ = 0.11, p = 0.0683; Pearson’s r = 0.11, p = 0.0561 (Fig. 3.d) indicating that methylation likely does not play a significant role in regulating *ATP2B2* expression in GBM. A weak negative association was observed between *ATP2B4* promoter methylation and its mRNA expression (Spearman’s r ≈ −0.05, p > 0.3; Pearson’s r ≈ −0.15, p = 0.0135). Although Pearson’s correlation reached statistical significance, the effect size was small indicating that promoter methylation is unlikely to be a major determinant of *ATP2B4* expression in this cohort (Fig. 3.e).

To further investigate the transcriptional pathways that regulate the simultaneous downregulation of neuroplastin and PMCA isoforms, we utilized the ChEA3 transcription factor (TF) enrichment platform to pinpoint non-methylation-associated upstream regulators. A clusterogram of TF-gene relationships across several datasets, including ENCODE, GTEx, ARCHS4, and curated literature, demonstrated significant overlap in transcription factor predictions for *NPTN* and *ATP2B4*. Notably, *ATP2B1* and *ATP2B2* exhibited a more modular pattern of TF-gene interactions. (Fig. 3.f). Complementing this matrix analysis, the ChEA3 TopRank output identified TFs with the highest integrated regulatory scores for this four-gene set. Among the top-ranked TFs were notably MBNL2 and ZEB2, factors implicated in the regulation of alternative splicing and epithelial-mesenchymal transition in cancer^50,51^ (Fig. 3.g). These transcription factors may function as upstream regulators of calcium dysregulation in GBM, thereby connecting transcriptional reprogramming to disrupted ion homeostasis.

### Protein-level downregulation of neuroplastin and PMCA in glioblastoma

To build on what we learned from transcriptional and epigenetic studies, we looked into whether the changed gene expression of Np and PMCA isoforms was reflected at the protein level (Fig. 4). In addition, we examined the protein expression of sodium/calcium exchanger 1 (NCX-1) to rule out the potential involvement of that calcium transporter in calcium dysregulation.

**Fig. 4.**
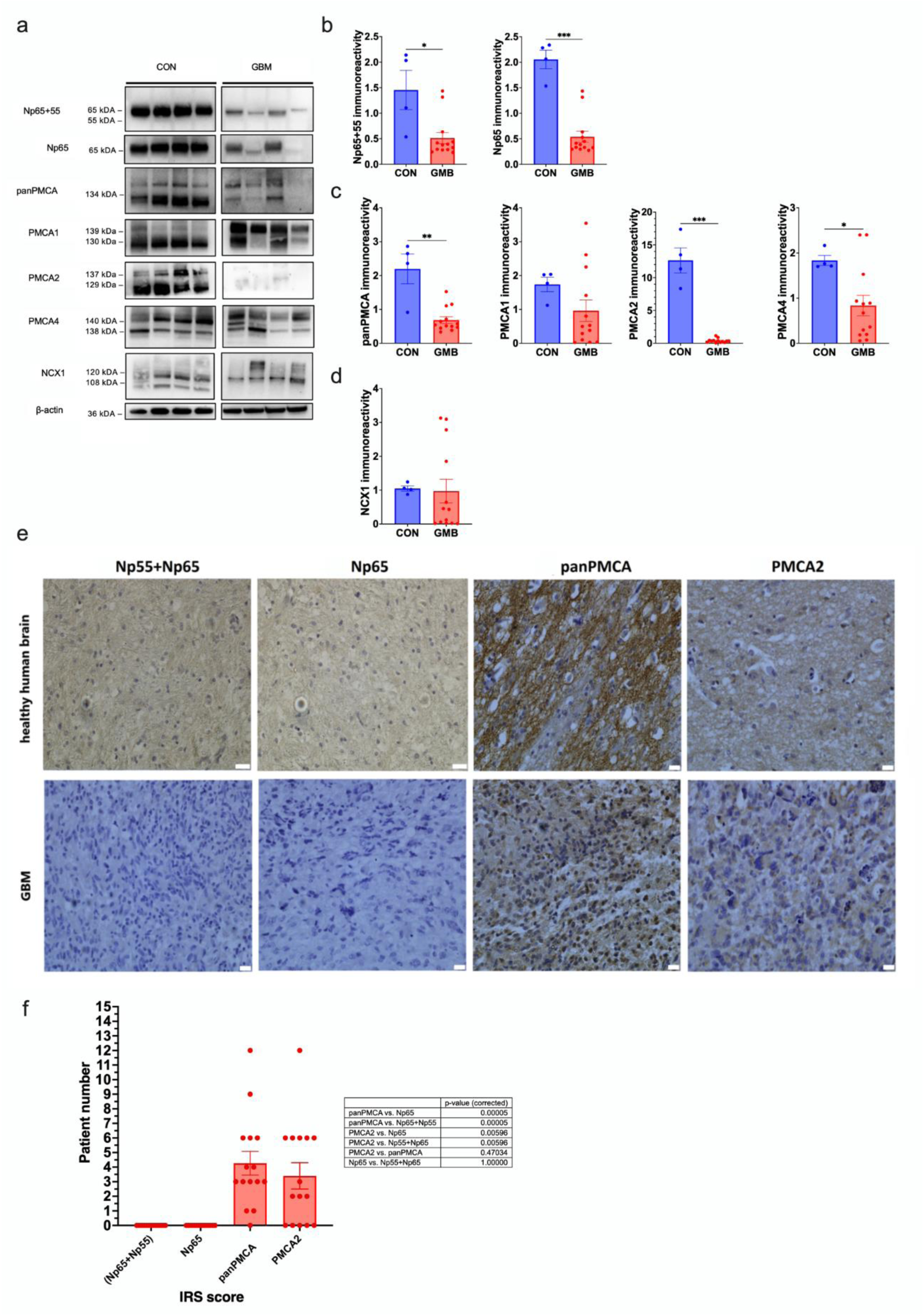
Protein expression analysis of Np, PMCA isoforms, and NCX-1 in control (CON) and glioblastoma (GBM) samples. a Representative Western blots showing Np65+Np55, Np65, pan-PMCA (total PMCA), PMCA1, PMCA2, PMCA4, and NCX-1 expression, with β-actin as the loading control. b Quantification of Np65+Np55 and Np65 protein levels. c Quantification of total PMCA (panPMCA) and PMCA isoforms. d NCX-1 expression quantification. All quantifications were normalized to total protein content per well, and data are presented as mean ± SEM, with individual data points overlaid. * = p < 0.05, ** = p < 0.001, *** = p < 0,0001; Man Whitney test. n = 4 CON; 13 GBM. e Immunohistochemical staining of healthy human brain tissue and glioblastoma for a) Np55+Np65, b) Np65, c) pan-PMCA, d) PMCA2. The figure shows tumor cells negative for expressing a) Np55+Np65 and b) Np65. Most cells in the tumor hot-spot area show low-to-moderate reactivity for c) PMCA and d) PMCA2 compared to the adjacent healthy brain tissue. Scale bar = 25 μm. f Immunoreactivity scores (IRS) of total neuroplastin (Np65+Np55), brain-specific Np65 isoform, PMCA2, and panPMCA with a table corresponding to pairwise *post-hoc* statistical comparisons of immunoreactive scores (IRS) amongst investigated proteins. *Post-hoc* analysis was conducted using the Mann-Whitney test with Bonferroni correction.

Western blot analysis showed that total neuroplastin (Np65+Np55) and brain-specific Np65 isoform protein levels were significantly reduced in GBM samples compared to controls (p = 0.0151, p = 0.008, respectively; Man-Whitney test; Fig. 4.b). Total PMCA (panPMCA), PMCA1, PMCA2, and PMCA4 protein levels were significantly decreased in GBM samples (p = 0.0059, p = 0.1630, p = 0.0008, p = 0.0445 respectively; Mann-Whitney test; Fig. 4.c). For the PMCA isoforms, PMCA1, PMCA2, and PMCA4, antibodies detect two distinct bands corresponding to each isoform’s splicing variants. In control samples, both PMCA1 variants were present at similar intensities. In contrast, in GBM samples, a reduction in both bands’ intensities was observed, with the b variant showing a more pronounced decrease (Fig. 4.a). PMCA2 exhibited a clear difference, with both PMCA variants being strongly detected in control samples. In contrast, in GBM samples, both bands were faint or nearly absent, with a greater reduction in the lower PMCA variant (Fig. 4.a). Similarly, PMCA4 showed a trend, with both PMCA4 variants exhibiting reduced intensity in GBM. NCX-1 expression was unchanged in GBM (p = 0.2958, Man-Whitney test), but showed significant variability, as indicated by the dispersion of individual values among all GBM samples (Fig. 4.d). The NCX-1 signal exhibited varied band patterns indicative of distinct molecular weights; control samples had several bands, with prominent signals at around 120 kDa and 108 kDa (Fig. 4.a). In GBM samples, the intensity of these bands was diminished and exhibited greater variability, with some samples showing weaker signals or the absence of certain bands, while others showed more prominent bands.

Immunohistochemical analysis of cellular protein expression revealed that the levels of the investigated proteins were generally very low or absent in our total GBM cohort compared with normal brain controls. The expression of four proteins, total neuroplastin (Np65 + Np55), the brain-specific Np65 isoform, PMCA2, and panPMCA, was quantified in a total of 15 glioblastoma samples using the immunoreactive score (IRS) method, which ranges from 0 to 12. All of the 15 glioblastomas showed a lack of expression or faintly detectable staining of both Np65 and Np65+Np55. The majority of cells counted in the tumor region show a lack of reactivity for Np65, Np65 + Np55, compared to positive controls (Fig. 4.e).

However, PMCA2 and pan-PMCA showed staining reactivity. PMCA2 with intensities of 0-1 in 60%, 2 in 33.3% and 3 in 6.7% of tumor cells. PanPMCA was stained with intensities of 0-1 in 47%, 2 in 40% and 3 in 13.3% (Fig. 4.f). PMCA2 and pan-PMCA expression levels were significantly higher than those of total neuroplastin and the brain-specific Np65 isoform, which were absent in all samples (IRS = 0). No significant difference was observed between PMCA2 and pan-PMCA, suggesting comparable levels of expression in the analyzed samples. A non-parametric Kruskal-Wallis test revealed a statistically significant difference in expression levels among the analyzed protein groups (H = 32.61, p = 3.89 × 10⁻⁷). Subsequent *post-hoc* analysis (Mann-Whitney test with Bonferroni correction) further delineated the pairwise differences, indicating that protein-level suppression was generally constant in GBM.

### Ganglioside expression patterns are altered in glioblastoma

In order to evaluate the aberrant ganglioside expression in GBM, we performed high-performance thin-layer chromatography (HPTLC) and dot blot analysis (Fig. 5).

**Fig. 5.**
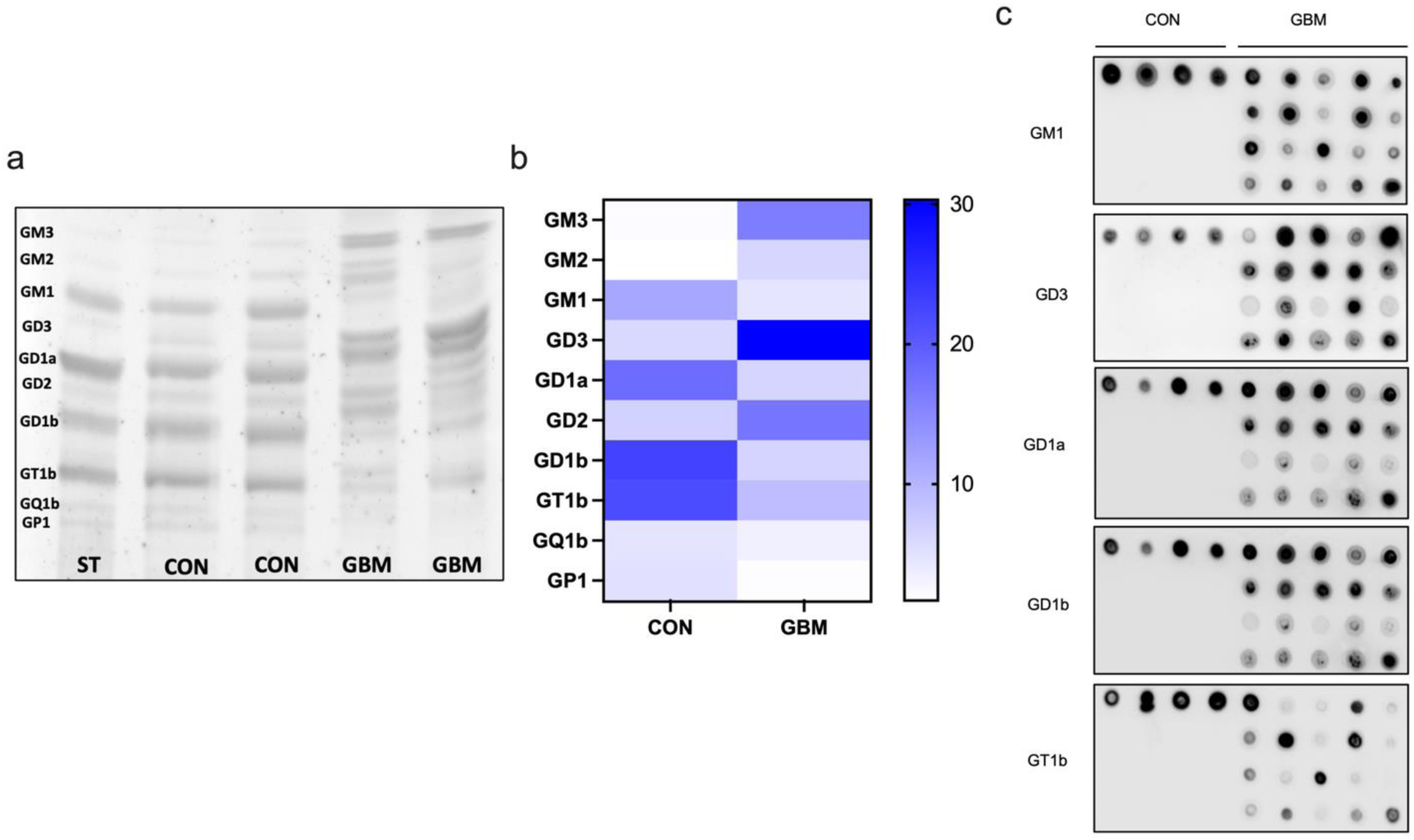
Analysis of ganglioside expression pattern in control (CON) and glioblastoma (GBM) samples. **a** HPTLC analysis showing the ganglioside composition of control and GBM samples. Gangliosides were separated based on their migration pattern, with bands corresponding to GM3, GM2, GM1, GD3, GD1a, GD2, GD1b, GT1b, GQ1b, and GP1; ST: standard. **b** Heatmap representation of ganglioside abundance in control and GBM samples, indicating relative intensity variations; n = 2/group. **c** Dot blot analysis of gangliosides in control (CON) and glioblastoma (GBM) samples. The figure illustrates the expression levels of GM1, GD1a, GD1b, GT1b, and GD3 gangliosides. Each ganglioside’s relative abundance is represented by dot intensity, with noticeable differences observed between control and GBM conditions.

HPTLC analysis showed that control brain tissue had a complex ganglioside profile that included GM3, GM2, GM1, GD3, GD1a, GD2, GD1b, GT1b, GQ1b, and GP1 (Fig. 5.a). In contrast, GBM samples exhibited a notable decrease in major brain gangliosides, particularly GM1, GD1a, GD1b, and GT1b. On the other hand, specific species, such as GD3, showed increased levels (Fig. 5.a). The accompanying heatmap further illustrates the differences in abundance of individual gangliosides according to quantification, with varying intensities observed for specific species (Fig. 5.b). In addition, dot blot analysis of gangliosides GM1, GD1a, GD1b, GT1b, and GD3 showed distinct differences in expression levels (Fig. 5.c), with dot intensity reflecting variations in the levels of specific gangliosides in glioblastoma samples compared to healthy brain controls. Complex gangliosides GM1, GD1a, GD1b and GT1b, in line with the results of HPTLC, showed overall lower signal intensities in GBM compared to controls; however signal intensity was highly variable across GBM samples, especially for GT1b (Fig. 5.c). In contrast, GD3 exhibited a notably higher signal in GBM than controls. These observations, in line with the literature, collectively demonstrate that GBM has a significantly modified ganglioside profile, characterized by a decrease in neuroprotective poly-sialogangliosides and an elevation in the simpler GD3 ganglioside.

### Integrated analysis reveals gangliosides as a key molecular determinants of reduced PMCA activity in glioblastoma

To evaluate whether PMCA function was also altered, in line with diminished mRNA and protein levels, we analysed PMCA activity. We further integrated those results into a principal component analysis (PCA) to potentially identify which factor has the most significant effect on altered PMCA activity (Fig. 6).

**Fig. 6.**
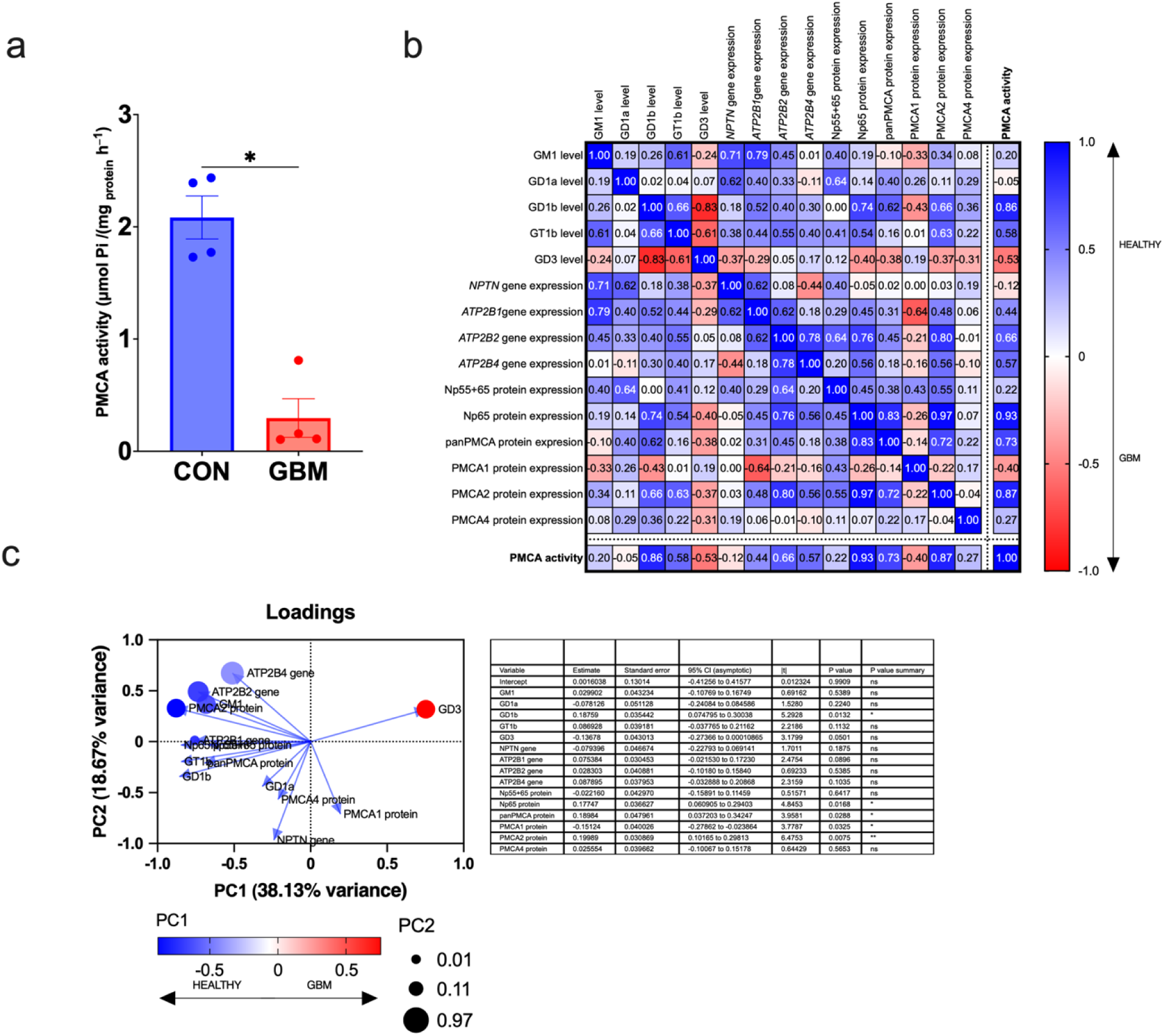
PMCA activity in glioblastoma (GBM) samples compared to controls (CON) and principal component analysis (PCA) investigating the factors mostly contributing to diminished PMCA activity. a Total PMCA activity was measured via inorganic phosphate (Pi) release from ATP hydrolysis in control (CON) and glioblastoma (GBM) samples. Activity is expressed as µmol Pi/h/mg protein. Data are shown as mean ± SEM. * = p < 0,05; Mann-Whitney test; n = 4 per group. b. Spearman correlation matrices constructed using z-score normalized values from the same cohort to explore relationships between ganglioside levels (GM1, GD1a, GD1b, GD3, GT1b), neuroplastin, and PMCA mRNA expression (*NPTN*, *ATP2B1*, *ATP2B2*, *ATP2B4*), and protein expression levels (Np65, Np65+Np55, PMCA1, PMCA2, PMCA2 and total PMCA (pan-PMCA)) with total PMCA activity across the sample set (n = 4 per group). Each matrix cell displays the Spearman correlation coefficient (ρ) between features. Red-highlighted correlations denote GBM-specific patterns, while blue indicates correlations preserved in controls. c. Principal Component Analysis (PCA) of z-scored molecular features related to PMCA regulation in the same cohort of GBM and control samples (n = 4 per group). The loading plot shows the contribution of individual variables to the first two principal components, with PC1 accounting for 38.13% and PC2 for 18.67% of the total variance. Features contributing the most to observed PMCA activity in GBM and control samples are highlighted in red and blue, respectively. The table indicates Principal Component Regression (PCR) coefficients for variables predicting PMCA activity in GBM and control samples.

Total PMCA activity was significantly reduced in GBM samples compared to control brain tissue (p = 0.0286, Mann-Whitney test), thus, tumor tissue showed an approximately 80% decrease in enzyme activity (Fig. 6.a). To unravel the molecular mechanisms underlying the severe diminishment of PMCA function, we analyzed gene expression, protein abundance, and ganglioside content in a cohort of four samples per group. Despite the limited sample size, the integrated data supported findings from large-scale datasets such as TCGA, enabling exploratory inference into key relationships. We conducted a descriptive Spearman correlation analysis, considering |ρ| > 0.7 as indicative of biologically meaningful associations (Fig. 6.b). Among gangliosides, GD1b showed the strongest positive correlation with PMCA activity in control samples (ρ = 0.86), suggesting a role in supporting PMCA function. Conversely, elevated GD3 levels were moderately associated with reduced PMCA activity in GBM (ρ = −0.53), though this did not reach statistical significance. At the protein level, Np65, PMCA2, and total PMCA (pan-PMCA) expression were all positively associated with the PMCA activity in control tissues (ρ = 0.93, 0.87, and 0.73, respectively). In GBM samples, lower PMCA1 expression showed the strongest negative correlation with PMCA activity (ρ = −0.40), although not statistically significant. Exploratory unsupervised principal component analysis (PCA) of z-scored molecular variables revealed that the first two principal components accounted for 38.1% and 18.7% of the total variance, respectively (Fig. 6.c). GD3 emerged as the primary contributor to variance along principal component analysis 1 (PCA1), underscoring its significance in the GBM molecular profile. Subsequently, principal component regression (PCR) analysis identified GD1b (p = 0.0132), Np65 protein (p = 0.0168), total PMCA protein (p = 0.0288), PMCA1 (p = 0.0325), and PMCA2 (p = 0.0075) as the most influential predictors of PMCA activity across all samples. These findings collectively emphasize the pivotal role of ganglioside composition, particularly GD1b and GD3, in modulating the coordinated calcium extrusion machinery involving neuroplastin and PMCA.

## Discussion

The data presented in this study reveals a disturbance of a previously unrecognized membrane-associated molecular nexus in GBM composed of specific isoforms of PMCA, Np, and gangliosides. Disruption of this molecular network in GBM impairs Ca²⁺ extrusion. Consequently, the inability to maintain normal intracellular Ca²⁺ levels can lead to intracellular Ca²⁺ overload that is associated with tumorigenesis.

Table 1. summarizes the gene and protein expression data from this study, comparing mRNA and protein levels with those from larger cohorts in publically available datasets, all in relation to specific ganglioside patterns in GBM. Our findings demonstrate that aberrant expression patterns of gangliosides (Fig. 5) in GBM, where complex gangliosides are downregulated, and tumorigenic gangliosides such as GD3 and GD2 are upregulated, go hand in hand with reduced Np expression, reduced isoform-specific PMCA expression, and impaired total PMCA activity.

**Table 1.**
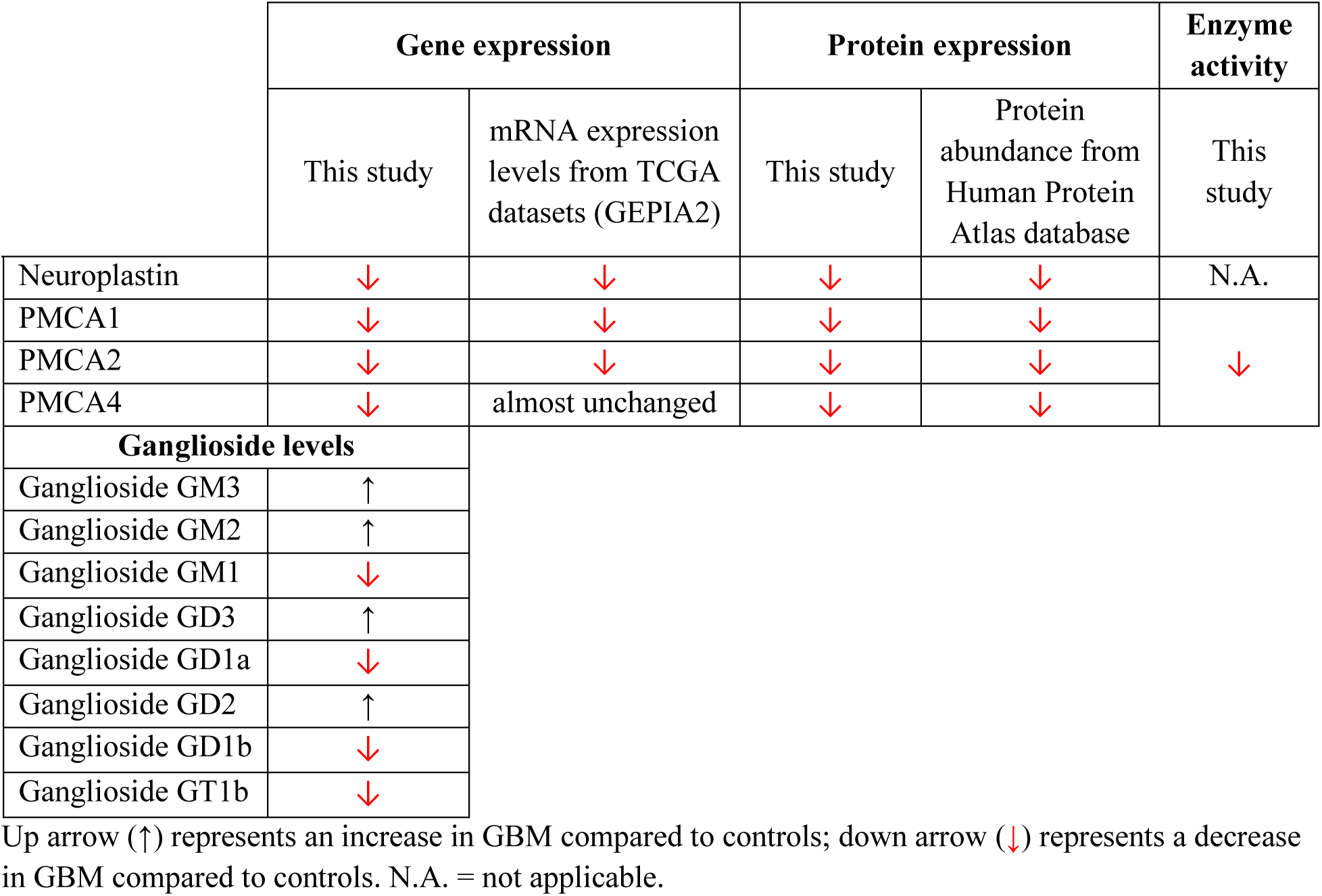
Summarized findings of the study.

|  | Gene expression |  | Protein expression |  | Enzyme activity |
| --- | --- | --- | --- | --- | --- |
|  | This study | mRNA expression levels from TCGA datasets (GEPIA2) | This study | Protein abundance from Human Protein Atlas database | This study |
| Neuroplastin | ↓ | ↓ | ↓ | ↓ | N.A. |
| PMCA1 | ↓ | ↓ | ↓ | ↓ | ↓ |
| PMCA2 | ↓ | ↓ | ↓ | ↓ |  |
| PMCA4 | ↓ | almost unchanged | ↓ | ↓ |  |
| Ganglioside levels |  |  |  |  |  |
| Ganglioside GM3 | ↑ |  |  |  |  |
| Ganglioside GM2 | ↑ |  |  |  |  |
| Ganglioside GM1 | ↓ |  |  |  |  |
| Ganglioside GD3 | ↑ |  |  |  |  |
| Ganglioside GD1a | ↓ |  |  |  |  |
| Ganglioside GD2 | ↑ |  |  |  |  |
| Ganglioside GD1b | ↓ |  |  |  |  |
| Ganglioside GT1b | ↓ |  |  |  |  |

Up arrow (↑) represents an increase in GBM compared to controls; down arrow (↓) represents a decrease in GBM compared to controls. N.A. = not applicable.

It has been established that the ganglioside pattern is markedly different in GBM compared to control tissues, regarding both concentration and composition. Gangliosides GD3 and GD2 are considered tumor markers, and the concentration of both has been found elevated in our study (Fig. 5), in accordance with the literature ^5,21–24^. Gangliosides are not mere inert architectural elements of the plasma membrane, but have an active role in signal transduction through direct interactions with specific proteins, as well as through providing an optimal microenvironment for the function of an array of proteins comprising the ganglioside interactome ^52^. Yet, the effects of particular gangliosides on membrane proteins’abundance and function in GBM are not being sufficiently investigated.

The proteins known to belong to the comprehensive ganglioside–interacting network are PMCA and Np. Gangliosides affect the activity of PMCA^37^, with poly-sialogangliosides such as GT1b having a stimulatory effect and mono-sialogangliosides such as GM3 having an inhibitory effect^25^. Since Np is an essential auxiliary subunit of PMCA, and the expression and microlocalization of Np is affected when the ganglioside composition of the membrane is changed ^37,53^, it is prudent to examine the expression of PMCA and Np in GBM together. Even more so since Np has been linked to various cancers (Supplementary Table 5), emerging as a tumor suppressor gene or facilitating tumorigenic processes depending on molecular context. In lung and breast cancer, Np overexpression is mainly associated with tumor promotion ^40,42–45^. To be more specific, Np was significantly related to asbestos-exposed lung tumors, where it was also shown that the gene was hypomethylated ^44^. The profiling of Np expression levels in lung cancer cell lines showed consistently high levels ^40^. In addition, a novel N*PTN-NRG1* fusion gene was identified in a case of advanced lung adenocarcinoma ^43^. Furthermore, an upregulation of *NPTN* mRNA was observed upon down-regulation of its regulatory long non-coding (lnc) RNA in non-small cell lung cancer ^47^. Results for breast cancer demonstrate that circulating tumor cells exhibit higher *NPTN* expression than breast cancer cell lines ^45^. *NPTN* was found to be highly expressed in approximately 20% of invasive breast carcinomas and 50% of breast carcinomas with distal metastasis using a breast cancer tissue array ^42^. On the other hand, downregulation of the *NPTN* gene in ITGB3 knockdown rat breast cancer cell clones has also been demonstrated ^46^. In adenomatous polyposis coli (APC)-driven tumorigenesis, *NPTN* downregulation indicates a tumor-suppressive role ^41^. Furthermore, a comprehensive RNA-seq analysis of gliomas in the OncoDB public database revealed decreased *NPTN* gene expression in tumors compared with normal tissue (Supplementary Figure 1). This is consistent with findings by Gong et al ^38^ who demonstrated that Np gene expression and Np protein were significantly decreased in higher-stage gliomas. The study was performed in paired glioma and control cohorts, identifying *NPTN* as a hub gene with altered chromatin openness and differential expression, serving as a novel cancer suppressor. Hence, the *in silico* analyses performed using data from publicly available databases are consistent with our experimental data, showing severe Np downregulation in GBM. Supplementary Table 5 provides an overview of studies investigating Np in various tumor types, underlining its potential importance in cancer biology. Since Np is expressed in different brain cells including neurons and glia (Supplementary Figure 2), it would be of interest to explore its’ expression in other tumor types as well. Following Np expression, individual PMCA isoforms were also found to be downregulated at both the gene and protein levels in our study (Table 1, Supplementary Tables 2-4, Supplementary Figures 4-6).

Furthemore, mutational burden analysis revealed frequent somatic mutations, with *ATP2B2* showing the highest rate (Supplementary Tables 2-4). Three independent prediction software programs identified several mutations as likely disease-causing, thereby further validating our key findings of lower gene expression in a smaller sample cohort and *in silico* across large datasets (TCGA, GTEx, COSMIC, and cBioPortal). Notably, Np65 and PMCA2 showed the most marked downregulation, suggesting that tumor evolution may be isoform-selective against the calcium-handling machinery of healthy neuronal tissue.

In silico analysis of TCGA GBM data with GEPIA2 indicated that elevated *NPTN* mRNA is associated with shorter overall and disease-free survival (Fig. 1.g). However, in our experimental cohort *NPTN* levels were uniformly low at both mRNA and protein levels. This discrepancy likely reflects differences between population-level bulk RNA datasets and and may be attributable to tumor subtype distribution, tumor cellularity, transcript isoform usage, or post-transcriptional regulation. Importantly, survival associations from GEPIA2 reflect median-based stratification across a heterogeneous cohort and do not imply that all GBMs express high *NPTN*.

In our study, we also examined the methylation status of the queried genes promoters to determine whether the lower mRNA expression is caused by epigenetic mechanisms (Fig. 3). For *NPTN* it was shown that its expression is largely methylation-independent. In contrast, *ATP2B1* displayed a moderate and significant positive correlation between methylation and mRNA expression implying that methylation supports its transcriptional activity. For *ATP2B2*, a non-significant positive correlation was detected indicating that methylation likely does not play a significant role in regulating *ATP2B2* expression in GBM. A weak negative association was observed between *ATP2B4* promoter methylation and its mRNA expression however, the effect size was small suggesting that methylation is not the primary regulator of *ATP2B4* expression in this cohort (Fig. 3.e). Collectively, these results indicate heterogeneous methylation-dependent transcriptional regulation among the analyzed genes, with *ATP2B1* showing a moderate methylation-associated increase in expression, *ATP2B4* exhibiting only a weak negative relationship indicative of limited promoter-driven repression, and both *NPTN* and *ATP2B2* remaining largely unaffected by methylation in GBM.

Other regulatory mechanisms, such as binding specific TFs including cancer-related ZEB2 and MBNL2 (Fig. 3), or histone modifications, may influence *ATP2B1* expression. Methylation’s weak positive correlation with *ATP2B2* expression also indicates it’s not a major regulator of *ATP2B2* gene expression in GBM and that other epigenetic factors or TFs may influence it. Gangliosides are known to interact with epigenetic regulators, such as histone deacetylases (HDACs) and DNA methyltransferases (DNMTs), thus leading to silencing of the *NPTN* or *PMCA* genes through either promoter methylation or chromatin remodeling ^54,55^. GD3, a key molecular determinant of reduced PMCA activity in GBM observed in our study (Fig. 6.c), was also found to confer resistance to p53-induced apoptosis in breast cancer by modulating mitochondrial function^56^. In addition, GD3 can activate transcription factors such as NF-κB, which regulates genes involved in inflammation, proliferation, and apoptosis ^57^. These findings exemplify that ganglioside composition alterations serve not only as passive markers of tumor phenotype but could actively modulate calcium homeostasis through the effect on gene transcription.

In addition to gene expression analyses, we evaluated Np and PMCA protein levels as well as PMCA activity (Fig. 4 and Fig. 6.a, Table 1). In the attempt to fully explore the effects of the specifically altered ganglioside environment in GBM (Fig. 5) we additionaly analysed Spearman correlation matrices constructed using z-score normalized values of all of the parameters analysed in this study, as well as principal component analysis (Fig. 6.b-c). These analyses help converge the complex relationship between our data, highlighting the potential most significant contributors to the outcome, which is altered PMCA activity. The intricate interrelationship among Np, PMCA and gangliosides is illustrated by several findings. In Np-deficient mouse models, PMCAs fail to localize or function correctly, causing elevated intracellular Ca²⁺ levels ^37^. This aligns with our observation of ∼80% loss of PMCA activity in our GBM samples (Fig. 6). Gangliosides, particularly GM1, stabilize Np-PMCA complexes in lipid rafts, which are crucial for neuronal calcium regulation^37^. Decreased expression of other major brain gangliosides, GD1a, GD1b, and GT1b (Fig. 5), could influence membrane microdomain organization and subsequent signaling cascades ^58^, with GD1a and GD1b known to interact with PMCAs, stabilizing their function and enhancing Ca²⁺ extrusion efficiency ^59,60^. Our small-scale Spearman correlation analysis has revealed that GD1b levels positively correlate with PMCA activity. In contrast, GD3 levels negatively associate with PMCA activity in GBM (Fig. 6.b-c), showing that an optimal ganglioside milieu is necessary for the proper functioning of this enzyme. This was further confirmed by PCA analysis; GD3 emerged as the dominant molecular feature attributed to GBM samples, while GD1b explained the variance in PMCA activity in controls (Fig. 6).

An unresolved question remains: whether the altered ganglioside composition initiates the cascade by disrupting the Np-PMCA complex at the membrane in specific membrane microdomains containing Np-PMCA-GM1 complex, or whether transcriptional repression of *NPTN* and *PMCA* genes by other ganglioside-causing mechanisms is the primary cause? The strong correlations between these genes and ganglioside biosynthetic genes (Fig. 2), along with the observed impact of ganglioside imbalance on PMCA activity (Fig. 6), suggest a complex, multilevel relationship in which the perturbation of ganglioside concentration and composition, together with protein repression, mutually reinforce, leading to further calcium dysregulation. Gangliosides not only affect the Np-PMCA complex but also modulate calcium signaling through other mechanisms including the effect on NCX1 ^61,62^. Although overall no change in NCX1 expression was observed in our study (Fig. 4.a-b), the marked cellular heterogeneity of GBM encompassing neural stem-like, astrocytic, oligodendrocyte progenitor-like, and neuronal-like cells potentially explains our finding. This diversity underscores the importance of accounting for intratumoral subpopulations when designing targeted therapies.

To conclude, our transcriptomic, epigenetic, proteomic, lipidomic, and *in silico* analyses consistently demonstrated downregulation of Np and PMCA isoforms both at mRNA and protein levels in our GBM cohort (Figs. 1, 4). These findings are consistent with a broader trend observed in public datasets (TCGA, GTEx, COSMIC, cBioPortal). Changes in gene and protein expression led to a ∼80% loss of PMCA activity in our GBM samples. Dysregulation involving the gangliosides-PMCA-Np triad offers several specific molecular targets for therapeutic development. Potential strategies include developing inhibitors of GD3 synthase or agents that normalize ganglioside composition to stabilize the Np-PMCA complex and restore calcium homeostasis. Gene therapy or small molecules that upregulate Np65 expression could also be considered, as they may rescue PMCA activity and reduce calcium dysregulation in neuronal-like tumor cells. Additionally, developing small molecules or peptides that mimic Np’s stabilizing effect on PMCA isoforms could counteract impaired pump function. Since GBM is uniformely lethal and the emerging evidence suggests that dysregulated calcium signaling plays a pivotal role in GBM’s proliferation, invasion, and treatment resistance^8–10,63,64^, we propose that an integrated approach encompassing PMCA, Np and gangliosides must be employed to understand GBM etiology and behaviour, and to develop new treatments aimed at improving the dire survival and recovery prognoses.

## Methods

### Human Samples

Glioblastoma samples graded 4 and FFPE slides of paired tumor tissues were collected from the University Hospital Center Zagreb and University Hospital Center Sestre Milosrdnice, Zagreb. A certified neuropathologist (A.J.) diagnosed CNS WHO grade 4 astrocytomas (IDH1 wt) in accordance with the latest WHO classification. The patients had no family history of brain tumors and were untreated before surgery, which could affect molecular analyses. The sample included 24 astrocytomas grade 4 (16 males, 8 females), aged 30 to 80 years (mean 58.75, median 51, standard deviation 14.437). The average age at diagnosis for males was 54 (median, 57; standard deviation, 12.579), while for females, it was 66 (median, 71; standard deviation, 14.585). Informed consent was obtained from the patients. The control group consisted of 4 post-mortem cerebrocortical tissue samples from individuals with no family history of brain tumors or other neuropathological conditions who died from cardiac arrest. Samples consisted of 3 males and one female, aged 69-84 (average 77, median 77.5, standard deviation 6.48). The study was approved by the School of Medicine Ethics Committee (case number: 380–59–10, 106-14-55/147; class: 641–01/14–02/01 and case number: 380-59-10106-17-100/134; class: 641-01/17-02/01), the University Hospital Center Sestre Milosrdnice (EP-7426/14-9, EP-5429/17-5) and the Ethics Committee of University Hospital Center Zagreb (number 02/21/JG, class: 8.1.-14/54–2).

### Quantitative Real-Time PCR (qRT-PCR)

Total RNA was isolated from brain tissue samples of healthy individuals and glioblastoma patients using the GeneJET RNA Purification Kit (Thermo Fisher Scientific, #K0702). Genomic DNA contamination was removed by DNase I treatment (Sigma-Aldrich, #EN0521), after which equal amounts of RNA were reverse-transcribed into complementary DNA (cDNA) using the High-Capacity cDNA Reverse Transcription Kit with RNase Inhibitor (Applied Biosystems, #4388950). qRT-PCR was performed using the SYBR Green PCR Master Mix (Applied Biosystems, #4309155) on a fixed quantity of cDNA. Primer sequences are detailed in Supplementary Table 6. The ΔΔCt method was applied for relative gene expression analysis, with β-actin as an endogenous normalization control. Fluorescence detection was conducted on a 7900 HT Real-Time PCR System (Applied Biosystems).

### Methylation-Specific PCR (MSP)

Genomic DNA was modified by bisulfite treatment using the MethylEdge™ Bisulfite Conversion System (Promega, Madison, WI, USA), following the manufacturer’s protocol. The bisulfite-converted DNA was then subjected to methylation-specific PCR (MSP) to determine the methylation status of the *NPTN* promoter region. Primer sets specific for either the methylated or unmethylated sequences of the bisulfite-converted *NPTN* promoter were designed and employed for MSP analysis. The primer sequences, annealing temperatures, and expected product sizes are listed in Supplementary Table 7. Each 25 µL PCR reaction contained: 1× TaKaRa EpiTaq PCR buffer (Mg²⁺-free), 2.5 mM MgCl₂, 0.3 mM dNTP mix, 20 pmol of each primer (Sigma-Aldrich, USA), 25 ng of bisulfite-converted DNA, and 1.5 U of TaKaRa EpiTaq HS DNA Polymerase (TaKaRa Bio, USA). PCR amplification was performed using the following thermal cycling conditions: initial denaturation at 95 °C for 5 minutes; 35 cycles of 95 °C for 30 seconds, primer-specific annealing temperature (see Supplementary Table 7) for 30 seconds, and 72 °C for 30 seconds; followed by a final extension at 72 °C for 7 minutes. PCR products were separated by electrophoresis on a 2% agarose gel stained with SYBR Safe DNA Gel Stain (Invitrogen, Thermo Fisher Scientific, USA) and visualized under UV illumination. Methylated human control DNA (Promega, USA) and unmethylated EpiTect control DNA (Qiagen, Germany) were included as positive controls, while nuclease-free water served as a negative control to detect potential contamination.

### Western Blot Analysis

Brain tissue samples from healthy controls and glioblastoma patients (n = 4 per group) were homogenized with a Potter-Elvehjem homogenizer in RIPA buffer (150 mM NaCl, 1% Triton X-100, 0.5% sodium deoxycholate, 0.1% SDS, 50 mM Tris, pH 8.0) supplemented with protease inhibitors. Homogenates were centrifuged at 12,000 × g for 20 minutes at 4°C, and the supernatants were collected. Protein concentrations were measured using the Pierce™ BCA Protein Assay Kit (Thermo Fisher Scientific, #23227). Equal amounts of protein (15 μg) were separated on 4-12% NuPAGE™ Bis-Tris Mini Gels (Invitrogen, #NP0323BOX) in NuPAGE™ MOPS SDS Running Buffer (Invitrogen, #NP0001) and transferred onto PVDF membranes with the Mini Blot Module (Invitrogen, #B1000). Membranes were blocked with 5% skim milk in PBST for 1 hour at room temperature, then incubated overnight at 4°C with primary antibodies (Supplementary Table 8). After washing, membranes were incubated with peroxidase-conjugated secondary antibodies (Supplementary Table 8) for 1 hour. Protein bands were visualized using SuperSignal™ West Femto Substrate (Thermo Fisher Scientific, #34095) on the ChemiDoc MP Imaging System (Bio-Rad). Band intensities were quantified with Image Lab 6.1 software (Bio-Rad) and normalized to β-actin expression.

### Immunohistochemistry

Immunohistochemistry (IHC) was performed to assess the presence and expression levels of selected proteins and to confirm the Western blot results. IHC staining was conducted on 4 μm-thick paraffin-embedded tissue sections (FFPE) mounted on silanized glass slides (DakoCytomation, Glostrup, Denmark). Sections were deparaffinized in xylene (two cycles of 10 minutes each) and rehydrated through a graded ethanol series (100%, 96%, and 70%), each step repeated twice for 3 minutes, followed by immersion in distilled water for 5 minutes. Antigen retrieval was performed by heating the sections twice for 15 minutes at 450 W and once for 25 minutes at 220 W in citrate buffer (prepared by dissolving 2.1 g of citric acid monohydrate in 1 L of distilled water, with the pH adjusted to 6.0 using 2 M NaOH). Endogenous peroxidase activity was blocked by incubating the sections with 3% hydrogen peroxide for 20 minutes in the dark at room temperature. To reduce nonspecific binding, the sections were treated with a serum-free, ready-to-use protein block (Agilent Technologies, USA) for 30 minutes at +4°C. Sections were then incubated overnight at +4°C with primary antibodies (Supplementary Table 8). Secondary antibodies (Supplementary Table 8) were applied for 1 hour at +4°C. DAB from the Dako REAL Envision detection system (Peroxidase/DAB, Rabbit/Mouse, HRP; Agilent Technologies, USA) was prepared according to the manufacturer’s instructions and incubated with the tissue samples for 45 seconds, followed by counterstaining with hematoxylin. The human cerebral cortex served as a positive control. Negative controls were processed using the same staining procedure but without primary antibodies. Staining evaluation was independently performed by two blinded observers, using an Olympus BX52 microscope (Olympus Life Science). At least 300 cells in the tumor hot spot area were counted at 200× magnification to determine cell numbers and protein expression levels. Protein expression was quantified using the immunoreactivity score (IRS), which showed the best correlation with computational image analysis.

IRS was calculated by multiplying the percentage of positive cells (PP score) by the staining intensity (SI score). PP score categories were: <1–20% positive cells = score 1; 20–50% = score 2; 50–85% = score 3; and >85% = score 4. SI was scored as follows: no or weak staining = score 1; moderate staining = score 2; strong staining = score 3. The IRS ranged from 1 to 12, based on the combination of PP and SI scores.

### Ganglioside Isolation and Analysis

For the high-performance thin-layer chromatography (HPTLC), gangliosides were extracted from brain tissues (n = 4 per group). Tissues were homogenized in ice-cold redistilled water and centrifuged at room temperature at 450 × g for 15 minutes. Lipids were extracted with chloroform/methanol (1:2, v/v) and separated with water and chloroform. The upper phases were collected, evaporated to dryness, and purified using Sephadex G-25 (Merck, #G2580) gel filtration. Purified gangliosides were separated using HPTLC plates (Sigma Aldrich, #1.05631.0001) and visualized with a resorcinol-HCl reagent. Images were scanned on the ChemiDoc MP Imaging System (Bio-Rad) and analyzed with ImageJ software (NIH) to determine the relative abundance of each ganglioside species. For the dot blot analysis, one microliter of tissue homogenates (n = 24) was spotted onto nitrocellulose membranes (Bio-Rad, #1620112) and air-dried. Membranes were blocked with 5% skim milk in PBST for 1 hour and incubated overnight at 4°C with ganglioside-specific primary antibodies (Supplementary Table 8). After washing, membranes were treated with peroxidase-conjugated secondary antibodies (Supplementary Table 8) and visualized using SuperSignal™ West Femto Substrate (Thermo Fisher Scientific, #34095) on the ChemiDoc MP Imaging System (Bio-Rad).

#### Plasma Membrane Ca²⁺-ATPase Activity Assay

PMCA enzyme activity was measured spectrophotometrically in control (CON) and glioblastoma (GBM) tissue homogenates (n = 4 per group) by monitoring the release of inorganic phosphate (Pi) from ATP hydrolysis. All procedures were performed at 4°C unless otherwise specified. The reaction mixture for ATPase activity consisted of 50 μg of protein, 30 mM Tris-HCl buffer (pH 7.4), 3 mM MgCl_2_, 100 mM NaCl, 20 mM KCl, and 3 mM ATP. 1 μM carboxyeosin was used as a PMCA-specific inhibitor to suppress the baseline activity of other ATPases. Before adding ATP, reaction mixtures were pre-incubated at 37°C for 10 minutes. The enzymatic reaction was allowed to proceed for 15 minutes and then stopped with 10% trichloroacetic acid (w/v) in an amount equal to the reaction volume. The samples were then centrifuged at 20,000 g for 10 minutes at 4°C, and the supernatant was collected for measuring the Pi amount. Equal volumes of chromogenic reagent, composed of 150 mM ferrous sulfate dissolved in 1% (w/v) ammonium heptamolybdate in 0.6 M sulfuric acid, were added. The absorbance of the resulting product was measured at 700 nm using a Varian Cary 100 Bio spectrophotometer (SpectraLab Scientific Inc.). The specific PMCA activity was calculated by subtracting the total ATPase activity in the absence of carboxyeosin from that in its presence. Results are expressed as µmoles of P_i_ released per hour per milligram of protein (μmol P_i_/h/mg protein). All reactions were performed in duplicate.

### *In silico* analysis of protein-protein interactions, expression patterns, mutational burden, methylation patterns, and survival on public databases, cBioPortal, COSMIC, and GEPIA

To assess the compatibility of our experimental findings, we analyzed publicly available data. Functional enrichment analysis of *NPTN*, *ATP2B1*, *ATP2B2*, and *ATP2B4* was performed using ShinyGO v0.85 (https://bioinformatics.sdstate.edu/go/), identifying overrepresented pathways related to ion homeostasis and signaling in the GO Biological Process database (FDR < 0.05). We used the STRING v11.5 database (https://string-db.org) with a medium confidence score threshold (0.4) to analyze protein-protein interactions between neuroplastin and PMCA 1, 2, and 4. We visualized the results to identify potential functional networks. We retrieved genetic alterations reported in the *NPTN*, *ATP2B1*, *ATP2B2*, and *ATP2B4* genes from the cBioPortal for Cancer Genomics (https://www.cbioportal.org) and the COSMIC databases ^65,66^. We accessed cBioPortal in September 2025. Transcription factor (TF) enrichment was conducted using the ChEA3 web-based platform (https://maayanlab.cloud/chea3/) to identify potential upstream regulators of *NPTN*, *ATP2B1*, *ATP2B2*, and *ATP2B4* genes using TopRank analysis. We also examined methylation patterns on cBioPortal. The Methylation Array Harmonization Workflow quantifies methylation levels at known CpG sites as beta values from raw methylation array data generated from multiple generations of Illumina Infinium DNA methylation arrays, including the HumanMethylation 27 (HM27) and HumanMethylation 450 (HM450) arrays. Survival plots were created using the Gene Expression Profiling Interactive Analysis (GEPIA2) database (http://gepia2.cancer-pku.cn) ^67^. Kaplan-Meier curves were generated and compared using the log-rank test. We performed survival analysis based on the expression status of the queried genes and plotted Kaplan-Meier curves. Pathogenicity of the reported mutations was assessed using PolyPhen-2 (http://genetics.bwh.harvard.edu/pph2/)^68^, SIFT (Sorting Intolerant from Tolerant algorithm) (https://sift.bii.a-star.edu.sg)^69^, and Mutation Taster (http://www.mutationtaster.org/)^70^ tools.

### Statistical analyses

All data are presented as mean ± standard error of the mean (SEM). As appropriate, group comparisons were conducted using the unpaired two-tailed *t*-tests or the Mann-Whitney tests based upon normality assessment by the Shapiro-Wilk test. A p-value < 0.05 was considered statistically significant. Sample sizes for experiments were estimated a priori with G*Power ^71^ based on effect sizes from preliminary enzyme activity data, and at least 3 samples per group (α = 0.0,9911288; Df = 4; critical t = 2,776445). All assays were performed in technical duplicates or triplicates, depending on sample availability.

Spearman’s rank correlation was used to assess relationships among transcripts, proteins, ganglioside levels, and enzymatic activities on z-score standardized data from a subset of samples from which all the analyses were performed. Exploratory principal component analysis (PCA) and principal component regression (PCR) were performed on the same data subset with PMCA enzyme activity as the outcome variable. PCA reduced and visualized the multivariate structure, while PCR used enzyme activity as the outcome and principal components as predictors. All statistical analyses and graphical presentations were performed using GraphPad Prism 9 (GraphPad Software Inc.).

## Data Availability

The authors declare that the data supporting the findings of this study, including experimental procedures and compound characterization, are available within the paper and its Supplementary Information files, or from the corresponding authors upon request.

## Acknowledgements

This work was supported by research projects funded by the Croatian Science Foundation (grants NeuroReact, IP-2016-06-8636 to S.K.B.), European Union through the European Regional Development Fund, Operational Programme Competitiveness and Cohesion (grant agreement No. KK.01.1.1.01.0007, CoRE – Neuro) and University of Zagreb research support grants NEURO-MOD-PUMP (10106-24-1546 to K.M.J.) and GLIOEMT (10106-24-1201 to N.P.Š) .

## Ethics declarations

The authors declare no competing interests.

## Contributions

B.P., K.M.J., and N.P.Š. conceived the study. B.P., A.K., N.M.H., A.B., V.P., F.D., A.K.V., A.U., E.J. and N.P.Š. performed molecular experiments and analyses. N.N. and A.J. provided patient samples, neuropathological evaluations, and clinical data. B.P., A.K., A.U. and N.P.Š. performed data visualization and figure generation. B.P., A.K., and N.P.Š. performed statistical and *in silico* analyses. S.K.B. and K.M.J. provided critical feedback on experimental design and interpretation. B.P., A.K., S.K.B., K.M.J., and N.P.Š. prepared the manuscript. K.M.J, S.K.B. and N.P.Š. provided the funding for the research. All authors have revised and approved the final version of the manuscript.

## Notes

### Competing Interest Statement

The authors have declared no competing interest.

### Summary of Updates

Supplementary material updated is now also attached.

