## Supplementary material for "Disruption of the ganglioside-plasma membrane calcium ATPase-neuroplastin nexus leads to impaired calcium homeostasis in glioblastoma"

**Supplementary Table 1.** Mutational spectrum of the *NPTN* gene in glioblastoma (GBM) from the COSMIC database

| Histology subtype 2 | AA mutations | AA mutations detailed | CDS mutations | CDS mutations detailed | Mutation taster | PolyPhen2 | SIFT | Sample ID | Pubmed ID |
|---|---|---|---|---|---|---|---|---|---|
| GBM | p.L231Q | Substitution - Missense, position 231, L➞Q | c.692T>A | Substitution, position 692, T➞A | Disease causing | Probably damaging | Affect protein function | 2955352 | 28916733 |
| GBM | p.L231Q | Substitution - Missense, position 231, L➞Q | c.692T>A | Substitution, position 692, T➞A | Disease causing | Probably damaging | Affect protein function | 2955353 | 28916733 |
| GBM | p.V212F | Substitution - Missense, position 212, V➞F | c.634G>T | Substitution, position 634, G➞T | Polymorphism | Probably damaging | Tolerated | 2491105 | 26496030 |
| GBM | p.E159D | Substitution - Missense, position 159, E➞D | c.477G>T | Substitution, position 477, G➞T | Polymorphism | Benign | Tolerated | 2963697 | 25642631 |
| GBM | p.R156H | Substitution - Missense, position 156, R➞H | c.467G>A | Substitution, position 467, G➞A | Disease causing | Benign | Tolerated | 2813454 | 28263318 |
| GBM | p.D152E | Substitution - Missense, position 152, D➞E | c.456C>A | Substitution, position 456, C➞A | Polymorphism | Benign | Tolerated | 2178128 | CGP study COSU329 |
| GBM | p.E99K | Substitution - Missense, position 99, E➞K | c.295G>A | Substitution, position 295, G➞A | Disease causing | Probably damaging | Tolerated | 2963694 | 25642631 |

| | | | | | | | | | |
|---|---|---|---|---|---|---|---|---|---|
| GBM | p.T78N | Substitution - Missense, position 78, T➞N | c.233C>A | Substitution, position 233, C➞A | Disease causing | Probably damaging | Tolerated | 2963695 | 25642631 |
| GBM | p.? | Unknown | c.92-5044G>A | Substitution - intronic | Polymorphism | Unknown | Unknown | 2813453 | 28263318 |
| GBM | p.? | Unknown | c.92-5044G>A | Substitution - intronic | Polymorphism | Unknown | Unknown | 2813454 | 28263318 |
| GBM | p.? | Unknown | c.92-5164G>A | Substitution - intronic | Polymorphism | Unknown | Unknown | 2813453 | 28263318 |
| GBM | p.? | Unknown | c.92-5164G>A | Substitution - intronic | Polymorphism | Unknown | Unknown | 2813454 | 28263318 |
| Not specified | p.? | Unknown | c.-53C>T | Substitution | Polymorphism | Unknown | Unknown | 2194185 | CGP study COSU545 |

This table summarizes *NPTN* gene mutations in glioblastoma multiforme (GBM) samples from the COSMIC database. 13 mutations were found, including 8 missense and 5 intronic substitutions of unknown consequence. The table lists AA and CDS changes, predicted pathogenicity, and associated PubMed and COSMIC identifiers. Mutation consequences are annotated in COSMIC (canonical transcript).

**Supplementary Table 2**. Distribution and predicted impact of *ATP2B1* (PMCA1) mutations in glioma samples from the COSMIC database

| *Histology Subtype 1* | *Histology subtype 2* | *AA mutations* | *AA mutations Detailed* | *CDS mutations* | *CDS mutations Detailed* | *Mutation taster* | *Polyphen2* | *SIFT* | *Sample ID* |
|---|---|---|---|---|---|---|---|---|---|
| Astrocytoma Grade IV | GBM | p.? | Unknown | C.*2393A>T | Substitution | Unknown | Unknown | Unknown | 2307432 |
| Astrocytoma Grade IV | GBM | p.? | Unknown | C.*1548A>G | Substitution | Unknown | Unknown | Unknown | 2491103 |
| Astrocytoma Grade IV | GBM | p.? | Unknown | C.*567del | Deletion | Unknown | Unknown | Unknown | 2491102 |
| Oligodendroglioma Grade III | Anaplastic | p.R1101W | Substitution - Missense, position 1101, R➞W | C.3301C>T | Substitution, position 3301, C➞T | Polymorphism | Probably damaging | Affect protein function | 2693018 |
| Astrocytoma Grade IV | GBM | p.N1005S | Substitution - Missense, position 1005, N➞S | C.3014A>G | Substitution, position 3014, A➞G | Disease causing | Benign | Tolerated | 2813453 |
| Astrocytoma Grade IV | GBM | p.N1005S | Substitution - Missense, position 1005, N➞S | C.3014A>G | Substitution, position 3014, A➞G | Disease causing | Benign | Tolerated | 2813454 |
| Astrocytoma Grade IV | GBM | p.E987K | Substitution - Missense, position 987, E➞K | C.2959G>A | Substitution, position 2959, G➞A | Polymorphism | Probably damaging | Affect protein function | 2633979 |

| | | | | | | | | | |
|---|---|---|---|---|---|---|---|---|---|
| Astrocytoma Grade IV | GBM | p.K883N | Substitution - Missense, position 883, K➞N | C.2649G>C | Substitution, position 2649, G➞C | Disease causing | Possibly damaging | Tolerated | 2748710 |
| Astrocytoma Grade IV | GBM | p.W844R | Substitution - Missense, position 844, W➞R | C.2530T>A | Substitution, position 2530, T➞A | Disease causing | Probably damaging | Affect protein function | 2963695 |
| Astrocytoma Grade IV | GBM | p.A817D | Substitution - Missense, position 817, A➞D | C.2450C>A | Substitution, position 2450, C➞A | Disease causing | Probably damaging | Affect protein function | 2963694 |
| Astrocytoma Grade IV | GBM | p.A804V | Substitution - Missense, position 804, A➞V | C.2411C>T | Substitution, position 2411, C➞T | Disease causing | Possibly damaging | Affect protein function | 2634009 |
| Astrocytoma Grade IV | GBM | p.R741K | Substitution - Missense, position 741, R➞K | C.2222G>A | Substitution, position 2222, G➞A | Polymorphism | Benign | Tolerated | 2961796 |
| Astrocytoma Grade IV | GBM | p.L734I | Substitution - Missense, position 734, L➞I | C.2200C>A | Substitution, position 2200, C➞A | Polymorphism | Benign | Tolerated | 2178192 |
| Astrocytoma Grade IV | GBM | p.G673S | Substitution - Missense, | C.2017G>A | Substitution, position 2017, G➞A | Polymorphism | Benign | Tolerated | 2633982 |

| | | | | | | | | | |
|---|---|---|---|---|---|---|---|---|---|
| | | | position 673, G➞S | | | | | | |
| Astrocytoma Grade IV | GBM | p.D631V | Substitution - Missense, position 631, D➞V | C.1892A>T | Substitution, position 1892, A➞T | Disease causing | Benign | Tolerated | 2963695 |
| Astrocytoma Grade IV | GBM | p.G619D | Substitution - Missense, position 619, G➞D | C.1856G>A | Substitution, position 1856, G➞A | Polymorphism | Probably damaging | Affect protein function | 2963695 |
| Astrocytoma Grade IV | GBM | p.Q283= | Substitution - coding silent | C.849A>G | Substitution, position 849, A➞G | Silent | Silent | Silent | 2120297 |
| Astrocytoma Grade IV | GBM | p.D174G | Substitution - Missense, position 174, D➞G | C.521A>G | Substitution, position 521, A➞G | Disease causing | Probably damaging | Affect protein function | 2963694 |
| Astrocytoma Grade IV | GBM | p.L168I | Substitution - Missense, position 168, L➞I | C.502T>A | Substitution, position 502, T➞A | Disease causing | Possibly damaging | Tolerated | 2072593 |
| Astrocytoma Grade IV | GBM | p.L168I | Substitution - Missense, position 168, L➞I | C.502T>A | Substitution, position 502, T➞A | Disease causing | Possibly damaging | Tolerated | 1312974 |

| | | | | | | | | | |
|---|---|---|---|---|---|---|---|---|---|
| Astrocytoma Grade IV | GBM | p.S141F | Substitution - Missense, position 141, S➞F | C.422C>T | Substitution, position 422, C➞T | Disease causing | Benign | Affect protein function (low confidence of this prediction) | 2963698 |
| Astrocytoma Grade IV | GBM | p.K883N | Substitution - Missense, position 883, K➞N | C.2649G>C | Substitution, position 2649, G➞C | Disease causing | Possibly damaging | Tolerated | 2748710 |

This table lists 22 glioma samples with *ATP2B1* mutations: 18 missense, 1 deletion, 1 synonymous substitution, and 2 unknown functional consequences. Functional predictions were from MutationTaster, PolyPhen-2, and SIFT.Sample and PubMed IDs are from the COSMIC database. AA and CDS mutation positions are relative to the canonical transcript.

**Supplementary Table 3.** Comprehensive list of *ATP2B2* (PMCA2) mutations identified in glioma samples from the COSMIC database

| *Histology subtype 1* | *Histology subtype 2* | *AA mutations* | *AA mutations Detailed* | *CDS mutations* | *CDS mutations Detailed* | *Mutation Tester* | *Polyphen2* | *SIFT* | *Sample ID* | *Pubmed ID* |
|---|---|---|---|---|---|---|---|---|---|---|
| Astrocytoma Grade IV | GBM | p.K1163R | Substitution - Missense, position 1163, K→R | C.3488A>G | Substitution, position 3488, A→G | Disease causing | | Affect protein function Low confidence of prediction | 2961716 | 37792958 |
| Astrocytoma Grade IV | GBM | p.K1163R | Substitution - Missense, position 1163, K→R | C.3488A>G | Substitution, position 3488, A→G | Disease causing | | Affect protein function Low confidence of prediction | 2961772 | 37792958 |
| Astrocytoma Grade IV | GBM | p.R1097= | Substitution - coding silent | C.3291C>T | Substitution, position 3291, C→T | Silent | Silent | Silent | 2813453 | 28263318 |
| Astrocytoma Grade IV | GBM | p.R1097= | Substitution - coding silent | C.3291C>T | Substitution, position 3291, C→T | Silent | Silent | Silent | 2813454 | 28263318 |
| Astrocytoma Grade IV | GBM | p.I1023V | Substitution - Missense, | C.3067A>G | Substitution, position 3067, A→G | Polymorphysm | | Affect protein function | 2963695 | 25642631 |

| | | | | | | | | | | |
|---|---|---|---|---|---|---|---|---|---|---|
| | | | position 1023, I➞V | | | | | | | |
| Astrocytoma Grade IV | GBM | p.A914G | Substitution - Missense, position 914, A➞G | C.2741C>G | Substitution, position 2741, C➞G | Polymorphysm | | | 2748700 | 29802247 |
| Astrocytoma Grade IV | GBM | p.T855= | Substitution - coding silent | C.2565G>A | Substitution, position 2565, G➞A | Polymorphysm Silent | Silent | Silent | 2813453 | 28263318 |
| Astrocytoma Grade IV | GBM | p.T855= | Substitution - coding silent | C.2565G>A | Substitution, position 2565, G➞A | Polymorphysm Silent | Silent | Silent | 2120260 | CGP study COSU329 |
| Astrocytoma Grade IV | GBM | p.T855= | Substitution - coding silent | C.2565G>A | Substitution, position 2565, G➞A | Polymorphysm Silent | Silent | Silent | 2107998 | 23917401 |
| Astrocytoma Grade IV | GBM | p.T855= | Substitution - coding silent | C.2565G>A | Substitution, position 2565, G➞A | Polymorphysm Silent | Silent | Silent | 2813454 | 28263318 |
| Astrocytoma Grade IV | GBM | p.R767W | Substitution - Missense, position 767, R➞W | C.2299C>T | Substitution, position 2299, C➞T | New base T equals old base T | Probably damaging | Affect protein function | 2108005 | 23917401 |
| Astrocytoma Grade IV | GBM | p.R767W | Substitution - Missense, position 767, R➞W | C.2299C>T | Substitution, position 2299, C➞T | New base T equals old base T | | | 1337883 | CGP study COSU329 |

| | | | | | | | | | | |
|---|---|---|---|---|---|---|---|---|---|---|
| Astrocytoma Grade IV | GBM | p.P738S | Substitution - Missense, position 738, P→S | C.2212C>T | Substitution, position 2212, C→T | Polymorphysm | | | 2664395 | 28852847 |
| Astrocytoma Grade IV | GBM | p.G679S | Substitution - Missense, position 679, G→S | C.2035G>A | Substitution, position 2035, G→A | Disease causing | | | 2961786 | 37792958 |
| Astrocytoma Grade IV | GBM | p.R604Q | Substitution - Missense, position 604, R→Q | C.1811G>A | Substitution, position 1811, G→A | Polymorphism | | | 2963697 | 25642631 |
| Astrocytoma Grade IV | GBM | p.S501= | Substitution - coding silent | C.1503C>T | Substitution, position 1503, C→T | Silent | Silent | Silent | 2107987 | 23917401 |
| Astrocytoma Grade IV | GBM | p.S501= | Substitution - coding silent | C.1503C>T | Substitution, position 1503, C→T | Silent | Silent | Silent | 1337860 | CGP study COSU329 |
| Astrocytoma Grade IV | GBM | p.D454N | Substitution - Missense, position 454, D→N | C.1360G>A | Substitution, position 1360, G→A | Disease causing | | | 2664382 | 28852847 |
| Astrocytoma Grade IV | GBM | p.R437C | Substitution - Missense, position 437, R→C | C.1309C>T | Substitution, position 1309, C→T | Disease causing | | | 1910247 | CGP study COSU329 |

| | | | | | | | | | | |
|---|---|---|---|---|---|---|---|---|---|---|
| Astrocytoma Grade IV | GBM | p.G413E | Substitution - Missense, position 413, G→E | C.1238G>A | Substitution, position 1238, G→A | Disease causing | | | 2813519 | 28263318 |
| Astrocytoma Grade IV | GBM | p.V390I | Substitution - Missense, position 390, V→I | C.1168G>A | Substitution, position 1168, G→A | Disease causing | | | 2813453 | 28263318 |
| Astrocytoma Grade IV | GBM | p.V390I | Substitution - Missense, position 390, V→I | C.1168G>A | Substitution, position 1168, G→A | Disease causing | | | 2813454 | 28263318 |
| Astrocytoma Grade IV | GBM | p.R437C | Substitution - Missense, position 437, R→C | C.1309C>T | Substitution, position 1309, C→T | Disease causing | | | 1910247 | CGP study COSU329 |
| Astrocytoma Grade IV | GBM | p.E142= | Substitution - coding silent | C.426G>A | Substitution, position 426, G→A | Silent | Silent | Silent | 2178158 | CGP study COSU329 |
| Astrocytoma Grade IV | GBM | p.D143N | Substitution - Missense, position 143, D→N | C.427G>A | Substitution, position 427, G→A | Disease causing | Benign | Tolerated | 2178158 | CGP study COSU329 |
| Astrocytoma Grade IV | GBM | p.A114V | Substitution - Missense, position 114, A→V | C.341C>T | Substitution, position 341, C→T | Disease causing | Probably damaging | Tolerated | 2120349 | CGP study COSU329 |

| | | | | | | | | | | |
|---|---|---|---|---|---|---|---|---|---|---|
| Astrocytoma Grade IV | GBM | p.L234V | Substitution - Missense, position 234, L→V | C.700C>G | Substitution, position 700, C→G | Not possible | Benign | Tolerated | 2664382 | 28852847 |
| Astrocytoma Grade IV | GBM | p.D454N | Substitution - Missense, position 454, D→N | C.1360G>A | Substitution, position 1360, G→A | Disease causing | | | 2664382 | 28852847 |
| Astrocytoma Grade IV | GBM | p.? | Unknown | C.907+2980C>G | Substitution - intronic | Unknown | Unknown | Unknown | 2664382 | 28852847 |
| Astrocytoma Grade IV | GBM | p.P738S | Substitution - Missense, position 738, P→S | C.2212C>T | Substitution, position 2212, C→T | Disease causing | | | 2664395 | 28852847 |
| Astrocytoma Grade IV | GBM | p.? | Unknown | C.-460+43526T>C | Substitution - intronic | Unknown | Unknown | Unknown | 2307394 | 24705251 |
| Astrocytoma Grade IV | GBM | p.A914G | Substitution - Missense, position 914, A→G | C.2741C>G | Substitution, position 2741, C→G | Disease causing | | | 2748700 | 29802247 |

The *ATP2B2* gene, encoding PMCA2, showed the highest mutation rate among PMCA isoforms. 32 glioma samples had ATP2B2 variants: 21 missense, 9 synonymous, and 2 intronic. Predicted functional consequences were based on MutationTaster, PolyPhen-2, and SIFT algorithms, considering only canonical transcript variants.

**Supplementary Table 4.** Summary of *ATP2B4* (PMCA4) gene mutations in glioma samples retrieved from the COSMIC database

| *Histology Subtype 1* | *Histology subtype 2* | *AA mutations* | *AA mutations Detailed* | *CDS mutations* | *CDS mutations Detailed* | *Mutation Tester* | *Polyphen 2* | *SIFT* | *Sample ID* | *Pubmed ID* |
|---|---|---|---|---|---|---|---|---|---|---|
| Astrocytoma Grade IV | GBM | p.S17I | Substitution - Missense, position 17, S➞I | C.50G>T | Substitution, position 50, G➞T | Polymorphism | Benign | Tolerated | 2963695 | 25642631 |
| Astrocytoma Grade IV | GBM | p.L29V | Substitution - Missense, position 29, L➞V | C.85C>G | Substitution, position 85, C➞G | New base G equals old base | Probably damaging | Tolerated | 2664382 | 28852847 |
| Astrocytoma Grade IV | GBM | p.T105= | Substitution - coding silent | C.315G>A | Substitution, position 315, G➞A | Silent | Silent | Silent | 2813454 | 28263318 |
| Oligodendroglioma Grade III | Anaplastic | p.P139T | Substitution - Missense, position 139, P➞T | C.415C>A | Substitution, position 415, C➞A | Disease causing | Benign | Tolerated | 2693087 | 28270234 |
| Astrocytoma Grade IV | GBM | p.R178P | Substitution - Missense, position 178, R➞P | C.533G>C | Substitution, position 533, G➞C | Disease causing | Probably damaging | Affect protein function | 2963695 | 25642631 |
| Oligodendroglioma Grade III | Anaplastic | p.S249F | Substitution - Missense, position 249, S➞F | C.746C>T | Substitution, position 746, C➞T | Disease causing | Possibly damaging | Affect protein function | 1745112 | 22072542 |

| | | | | | | | | | | |
|---|---|---|---|---|---|---|---|---|---|---|
| Astrocytoma Grade IV | GBM | p.L380I | Substitution - Missense, position 380, L➞I | C.1138C>A | Substitution, position 1138, C➞A | Disease causing | Probably damaging | Tolerated | 2813453 | 28263318 |
| Astrocytoma Grade IV | GBM | p.L380I | Substitution - Missense, position 380, L➞I | C.1138C>A | Substitution, position 1138, C➞A | Disease causing | Probably damaging | Tolerated | 2813454 | 28263318 |
| Astrocytoma Grade IV | GBM | p.R488H | Substitution - Missense, position 488, R➞H | C.1463G>A | Substitution, position 1463, G➞A | Polymorphysm | Benign | Affect protein function | 2963697 | 25642631 |
| Astrocytoma Grade IV | GBM | p.L503F | Substitution - Missense, position 503, L➞F | C.1507C>T | Substitution, position 1507, C➞T | Disease causing | Probably damaging | Tolerated | 2955346 | 28916733 |
| Astrocytoma Grade IV | GBM | p.L503F | Substitution - Missense, position 503, L➞F | C.1507C>T | Substitution, position 1507, C➞T | Disease causing | Probably damaging | Tolerated | 2955350 | 28916733 |
| Astrocytoma Grade IV | GBM | p.L503F | Substitution - Missense, position 503, L➞F | C.1507C>T | Substitution, position 1507, C➞T | Disease causing | Probably damaging | Tolerated | 2955349 | 28916733 |
| Astrocytoma Grade IV | GBM | p.L503F | Substitution - Missense, | C.1507C>T | Substitution, position 1507, C➞T | Disease causing | Probably damaging | Tolerated | 2955348 | 28916733 |

| | | | | | | | | | | |
|---|---|---|---|---|---|---|---|---|---|---|
| | | | position 503, L➞F | | | | | | | |
| Astrocytoma Grade IV | GBM | p.L503F | Substitution - Missense, position 503, L➞F | C.1507C>T | Substitution, position 1507, C➞T | Disease causing | Probably damaging | Tolerated | 2955347 | 28916733 |
| Astrocytoma Grade IV | GBM | p.L546= | Substitution - coding silent | C.1636C>T | Substitution, position 1636, C➞T | Silent | Silent | Silent | 2178158 | CGP study COSU329 |
| Astrocytoma Grade IV | GBM | p.Q551* | Substitution - Nonsense | C.1651C>T | Substitution, position 1651, C➞T | Nonsense | Nonsense | Nonsense | 2178158 | CGP study COSU329 |
| Oligodendroglioma Grade II | Not specified | p.K561N | Substitution - Missense, position 561, K➞N | C.1683G>T | Substitution, position 1683, G➞T | Disease causing | Possibly damaging | Tolerated | 1745099 | 22072542 |
| Astrocytoma Grade IV | GBM | p.V578= | Substitution - coding silent | C.1734C>T | Substitution, position 1734, C➞T | Silent | Silent | Silent | 2178158 | CGP study COSU329 |
| Astrocytoma Grade IV | GBM | p.I579= | Substitution - coding silent | C.1737C>T | Substitution, position 1737, C➞T | Silent | Silent | Silent | 2178158 | CGP study COSU329 |

| | | | | | | | | | | |
|---|---|---|---|---|---|---|---|---|---|---|
| Astrocytoma Grade IV | GBM | p.F586= | Substitution - coding silent | C.1758C>T | Substitution, position 1758, C→T | Silent | Silent | Silent | 2178158 | CGP study COSU329 |
| Astrocytoma Grade IV | GBM | p.R587H | Substitution - Missense, position 587, R→H | C.1760G>A | Substitution, position 1760, G→A | Disease causing | Benign | Tolerated | 2963697 | 25642631 |
| Astrocytoma Grade IV | GBM | p.I596= | Substitution - coding silent | C.1788C>T | Substitution, position 1788, C→T | Silent | Silent | Silent | 2178158 | CGP study COSU329 |
| Astrocytoma Grade IV | GBM | p.E741* | Substitution - Nonsense | C.2221G>T | Substitution, position 2221, G→T | Nonsense | Nonsense | Nonsense | 2178192 | CGP study COSU329 |
| Astrocytoma Grade IV | GBM | p.A805T | Substitution - Missense,position 805, A→T | C.2413G>A | Substitution, position 2413, G→A | ? | Probably damaging | Tolerated | 2813454 | 28263318 |
| Astrocytoma Grade IV | GBM | p.? | Unknown | C.3309+70A>G | Substitution - intronic | Unknown | Unknown | Unknown | 2178075 | CGP study COSU329 |
| Astrocytoma Grade IV | GBM | p.? | Unknown | C.3309+70A>G | Substitution - intronic | Unknown | Unknown | Unknown | 2178075 | CGP study COSU329 |

| | | | | | | | | | | |
|---|---|---|---|---|---|---|---|---|---|---|
| Astrocytoma Grade IV | GBM | p.L29V | Substitution - Missense, position 29, L➞V | C.85C>G | Substitution, position 85, C➞G | New base G equals old base | Probably damaging | Tolerated | 2664382 | 28852847 |

Twenty-seven glioma samples had *ATP2B4* mutations, including 17 missense, 6 synonymous, 2 nonsense, and 2 intronic substitutions. MutationTaster, PolyPhen-2, and SIFT predicted functional impact. Mutation identifiers correspond to COSMIC sample IDs and PubMed references.

**Supplementary Table 5.** Overview of studies of neuroplastin expression in various cancers

| Cancer type | Type of sample | Sample size | Method of detection | Neuroplastin expression trend/major findings | Reference | Reference in the main text |
|---|---|---|---|---|---|---|
| Glioma | Database search with paired glioma and adjacent normal samples | 3 paired cohorts | Full-length RNA sequencing | ↓ *NPTN* gene expression | 1 | 38 |
| Pancreatic Cancer | Pancreatic cancer cell line BxPC-3 | N/A | Cell-surface glycoprotein capture | Neuroplastin identified | 2 | 39 |
| Lung cancer | Human lung cancer cell line A549 (KRAS-mutant: G12S; ATCC) | N/A | Immunocitochemistry | ↑ Neuroplastin protein expression induces cancerous events like motility and invasiveness | 3 | 40 |
| Colorectal cancer | Colorectal adenomatous polyps and | 42 human specimens | Microarray Expression Profiling | ↓ NPTN gene expression | 4 | 41 |
| Breast cancer | MDA-MB-435S cell line and primary breast tumors | 16 primary breast tumors | Transcriptomics | ↑ *NPTN* gene expression | 5 | 42 |
| Lung cancer | Lung cancer tissue | 1 human sample | Next generation sequencing | Novel *NPTN-NRG1* fusion identified in lung adenocarcinoma | 6 | 43 |

| | | | | | | |
|---|---|---|---|---|---|---|
| Lung cancer | Lung cancer samples associated with asbestos exposure | 91 sample pairs | Pyrosequencing | *NPTN* gene hypomethylated | 7 | 44 |
| Breast cancer | Circulating tumor cells (CTCs) in comparison to MCF7, SKBR3, T47D and MDA-MB-231 breast cancer cell lines | N/A | Microfluidic Dynamic Arrays | ↑ *NPTN* gene expression | 8 | 45 |
| Breast cancer | MDA-MB-231 breast cancer cell clones with conditional ITGB3 knockdown (I3 and I5) | N/A | Microarray analysis | ↓ *NPTN* gene expression | 9 | 46 |
| Non-small cell lung cancer | PC9-GR and PC9 non-small cell lung cancer cell lines | N/A | Transcriptomics | ↓ *NPTN* gene expression in the PC9-GR cell line | 10 | 47 |

*NPTN*: neuroplastin gene; ↑: higher expression compared to controls**;** ↓: lower expression compared to controls; N/A: not stated.

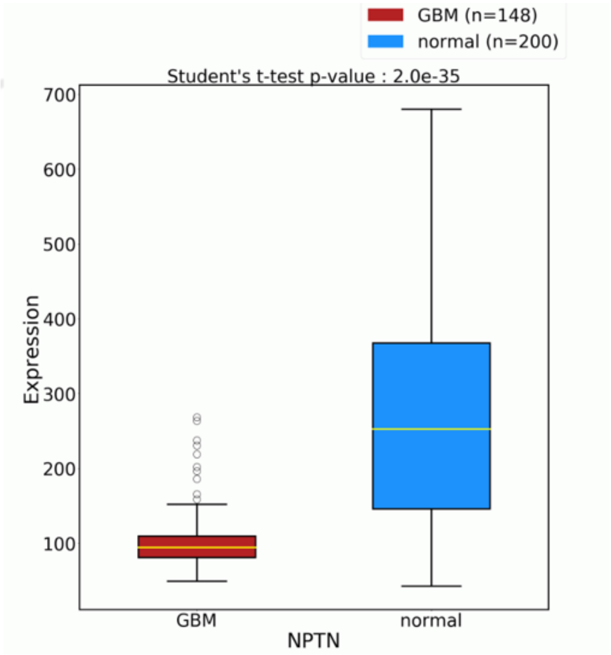


**Supplementary Figure 1**. Neuroplastin gene expression in glioblastoma as observed by RNA-Seq at the OncoDB public database. The data is normalized using TPM [11,12].

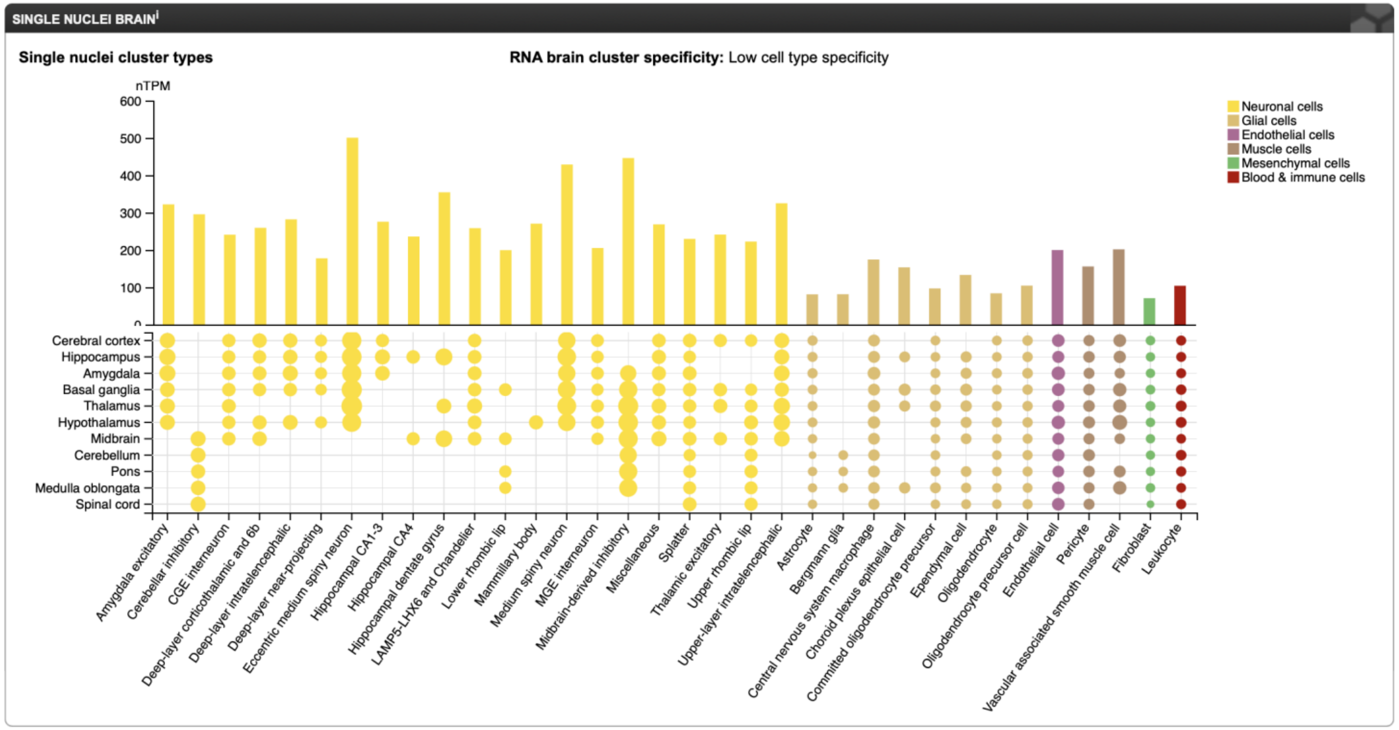


**Supplementary Figure 2**. Neuroplastin expression among different brain regions and cell types from Human Protein Atlas proteinatlas.org. This summary presents the normalized single nuclei RNA (nTPM) data derived from brain single nuclei in the Human Protein Atlas. The color coding reflects various cell type groups outlined in the single cell type data across the entire body, categorized into primary cell groups: neurons, glial cells, endothelial cells, fibroblasts, and muscle cells (including vascular smooth muscle cells and pericytes). The data is based on public data [13].

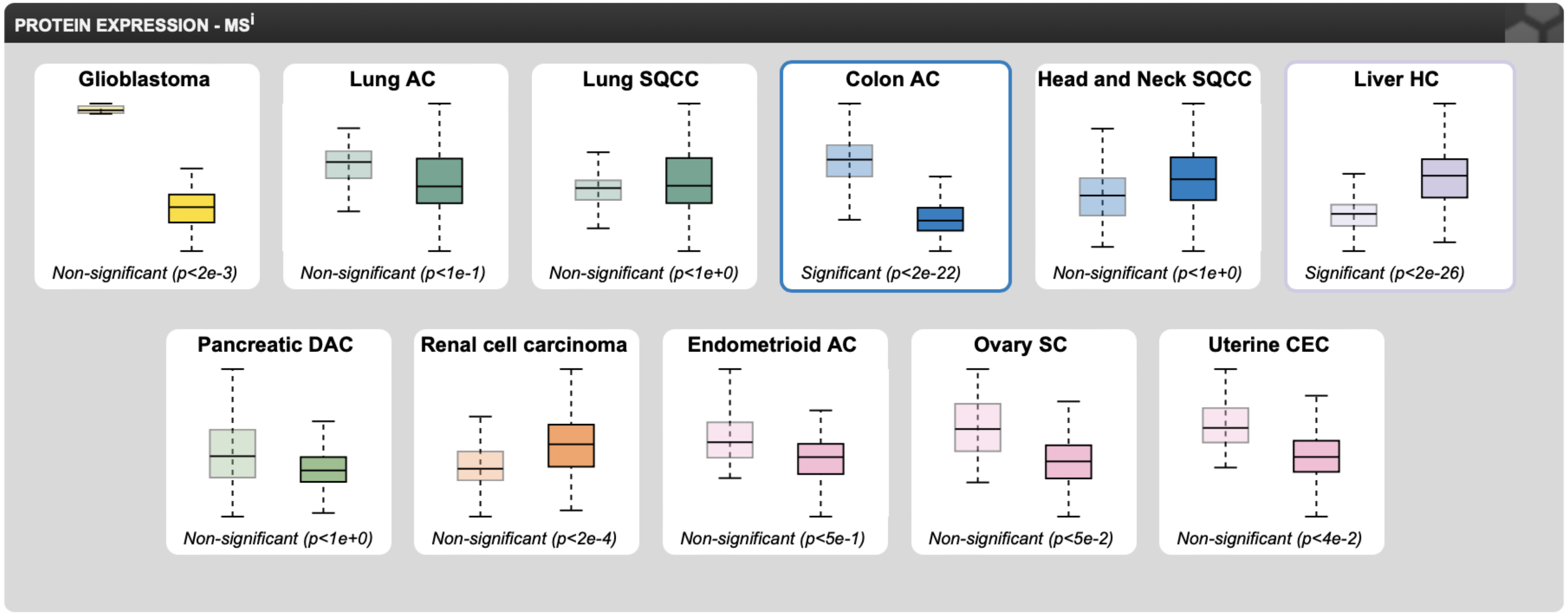


**Supplementary Figure 3**. Neuroplastin expression among different cancer types from Human Protein Atlas proteinatlas.org. nRPX is calculated as log2(intensity), derived from global protein abundance measurements at the protein level. These measurements are obtained through mass spectrometry experiments performed by CPTAC on tissues using isobaric tandem mass tags (TMT). On all images the left boxed plot represent normal tissue, while right boxed plot represents tumor tissue.

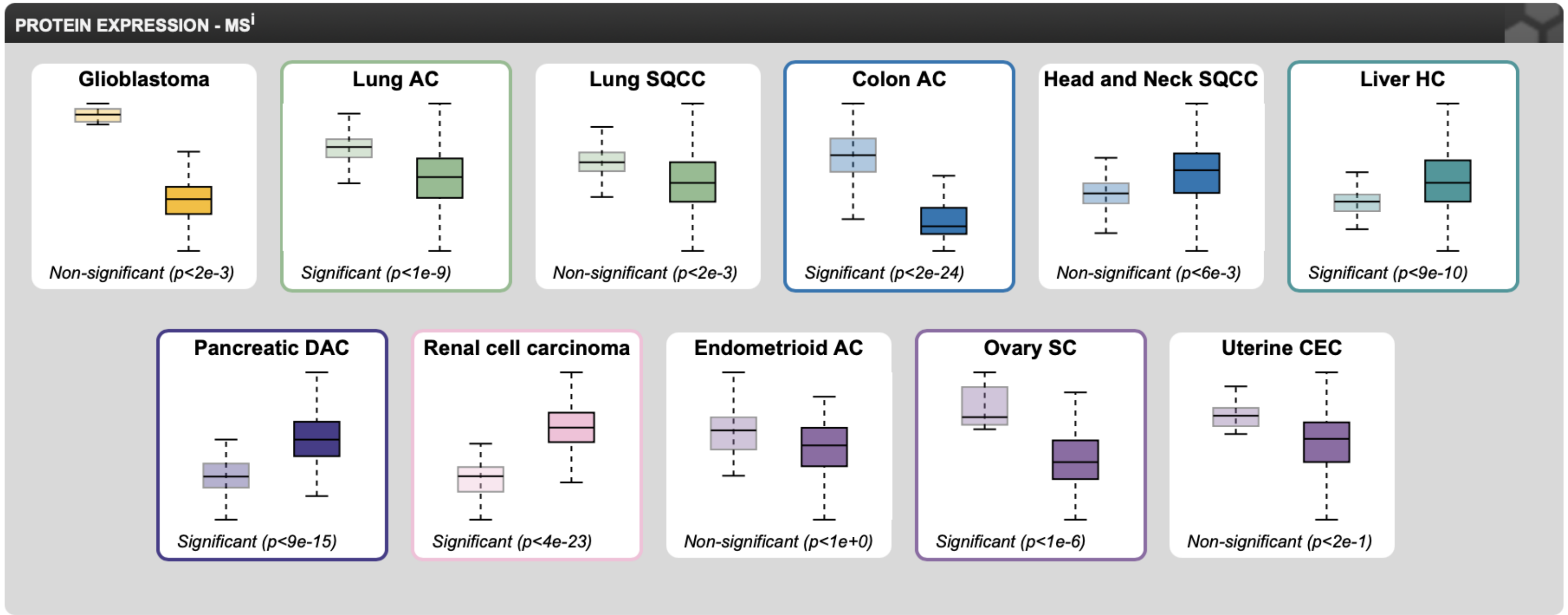


**Supplementary Figure 4**. PMCA1 expression among different cancer types from Human Protein Atlas proteinatlas.org. nRPX is calculated as log2(intensity), derived from global protein abundance measurements at the protein level. These measurements are obtained through mass spectrometry experiments performed by CPTAC on tissues using isobaric tandem mass tags (TMT). On all images the left boxed plot represent normal tissue, while right boxed plot represents tumor tissue.

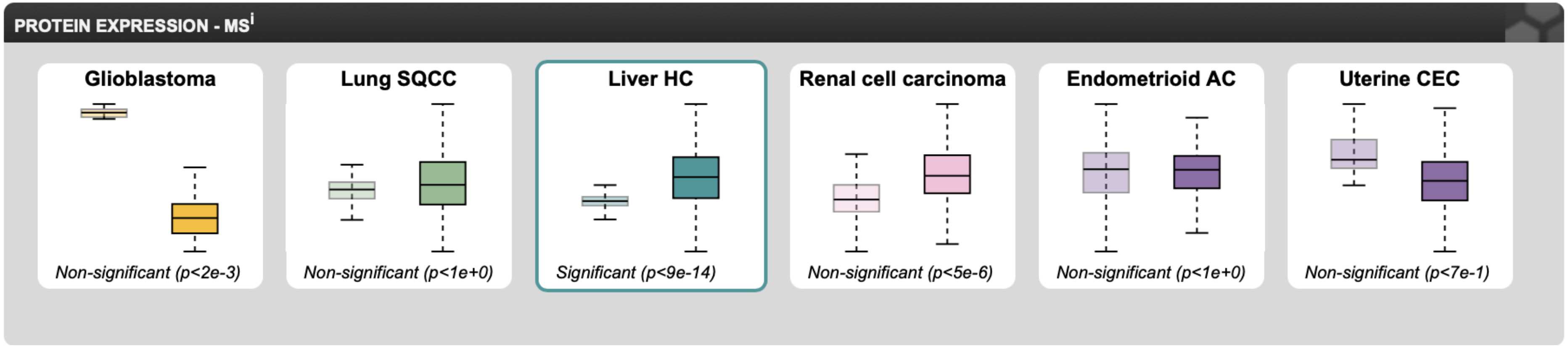


**Supplementary Figure 5**. PMCA2 expression among different cancer types from Human Protein Atlas proteinatlas.org. nRPX is calculated as log2(intensity), derived from global protein abundance measurements at the protein level. These measurements are obtained through mass spectrometry experiments performed by CPTAC on tissues using isobaric tandem mass tags (TMT). On all images the left boxed plot represent normal tissue, while right boxed plot represents tumor tissue.

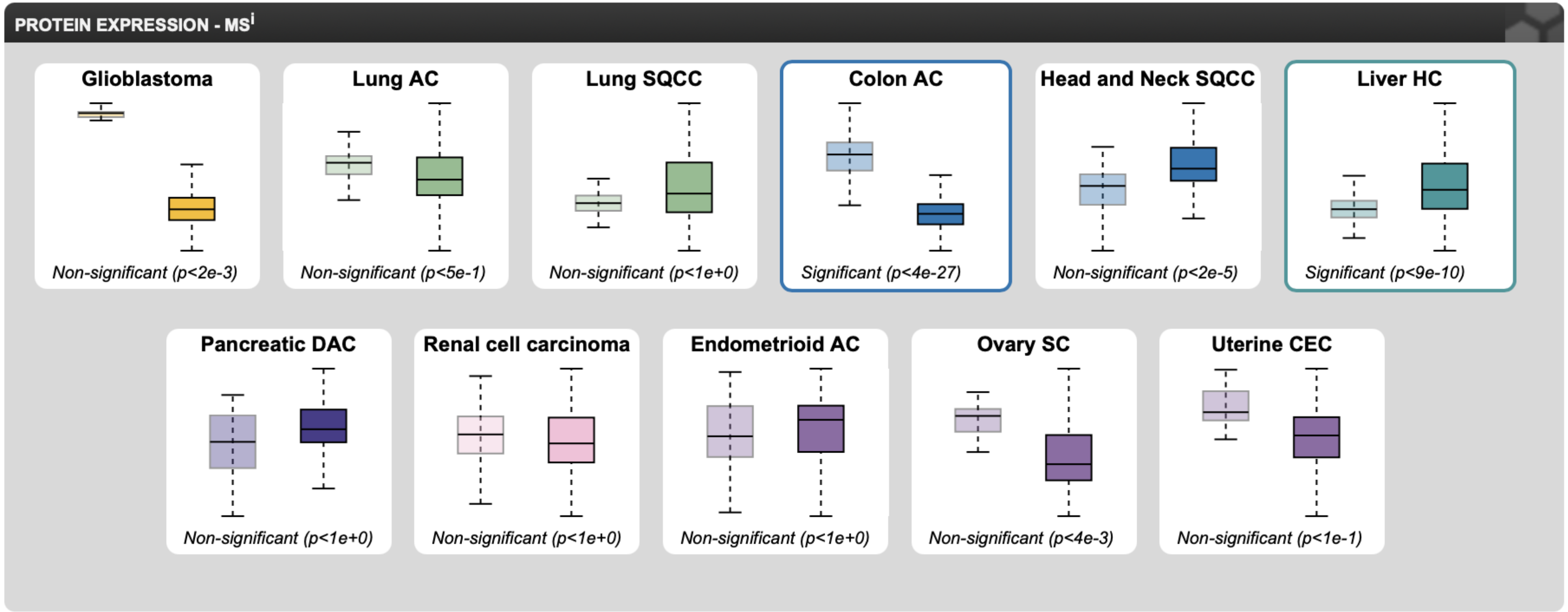


**Supplementary Figure 6**. PMCA4 expression among different cancer types from Human Protein Atlas proteinatlas.org. nRPX is calculated as log2(intensity), derived from global protein abundance measurements at the protein level. These measurements are obtained through mass spectrometry experiments performed by CPTAC on tissues using isobaric tandem mass tags (TMT). On all images the left boxed plot represent normal tissue, while right boxed plot represents tumor tissue.

**Supplementary Table 6.** qRT-PCR primer sequences given in 5'-3' direction

| Gene | Forward primer | Reverse primer | Reference |
|---|---|---|---|
| *NPTN* | ACCTGCAGATCACGGAAGAC | AGGACAGTGACAACAGAGGC | *In house* |
| *ATP2B1* | TTCAACGAAATAAATGCCCGG | AGGGTGGAGGACTGGAGTTACG | [14] |
| *ATP2B2* | ACAGTGGTACAGGCCTATGTCG | CGAGTTCTGCTTGAGCGCGG | [14] |
| *ATP2B4* | CAACTCCCGAAAGATCCATG | TGGTTAACACAGCAGCTGAC | [14] |
| *SLC8A1* | ACCACCAAGACTACAGTGCG | TTGGAAGCTGGTCTGTCTCC | [15] |
| *ACTB* | GAAGAGCTACGAGCTGCCTGA | CCACGTCACACTTCATGATGG | [16] |

**Supplementary Table 7.** Primer sequences used for methylation-specific PCR (MSP) analysis of the *NPTN* promoter region

| Primer | | Sequence | Annealing temperature | Product size (bp) |
|---|---|---|---|---|
| *NPTN* | MR - F | 5'- TTAGGTTAATTAGTTTTTGTAGGGGC -3' | 59,5 ℃ | 212 |
| | MR - R | 5'- AAAAATAAACGAAAAAAACAACCG -3' | | |
| | UMR - F | 5'- AGGTTAATTAGTTTTTGTAGGGGTG -3' | 58 ℃ | 211 |
| | UMR - R | 5'- AAAAAATAAACAAAAAAAACAACCAC -3' | | |

Primers differentiate methylated (MR) and unmethylated (UMR) DNA sequences after bisulfite conversion. Optimized annealing temperatures are reported for each primer pair. Amplicon sizes are in base pairs (bp).

**Supplementary Table 8.** List of primary and secondary antibodies used in Western blot and immunohistochemical analyses

| Antibody | Western blot dillution | Immunohistochemistry dillution | Dot blotting dilution | Supplier/ID |
|---|---|---|---|---|
| **Primary antibodies** | | | | |
| Rabbit polyclonal anti-neuroplastin 55+65 | 1:1000 | 1:500 | | *Donated by the courtesy of R.H.Mollina, Leibniz Institute for Neurobiology, Germany Prepared as in* [17] |
| Goat polyclonal anti-neuroplastin 65 | 1:1000 | 1:150 | | R&Dsystems, Minneapolis, MN, USA /AF5360 |
| Mouse monoclonal anti-panPMCA | 1:2000 | 1:2000 | | Santa Cruz/sc-271193 |
| Rabbit monoclonal anti-PMCA1 | 1:1000 | - | | Abcam, Cambridge, UK/ab190355 |
| Rabbit polyclonal anti-PMCA2 | 1:1000 | 1:200 | | Abcam, Cambridge, UK/ab3529 |
| Mouse monoclonal anti-PMCA4 | 1:1000 | - | | Abcam, Cambridge, UK/ab2783 |
| Rabbit monoclonal anti-NCX1 | 1:1000 | - | | Abcam, Cambridge, UK/ab177952 |
| Mouse monoclonal anti-β-Actin | 1:1000 | - | | Abcam, Cambridge, UK/ab8226 |
| Rabbit Polyclonal anti- beta subunit Cholera Toxin | | | 1:50 000 | Abcam, Cambridge, UK/ ab34992 |
| Mouse monoclonal anti GD1a | | | 1:2000 | *Donated by the courtesy of R.L. Schnaar, John Hopkins University; Baltimore, MA, USA;* 1.74 mg/mL |
| Mouse monoclonal anti GD1b | | | 1:2000 | *Donated by the courtesy of R.L. Schnaar, John Hopkins University; Baltimore, MA, USA;* 1.64 mg/mL |
| Mouse monoclonL anti-GT1b | | | 1:2000 | *Donated by the courtesy of R.L. Schnaar, John Hopkins University; Baltimore, MA, USA;* 2.44 mg/mL |
| Mouse monoclonL anti-GD3 | | | 1:250 | Abcam, Cambridge, UK/ ab11779 |
| **Secondary antibodies** | | | | |
| Donkey Anti-Mouse IgG HRP | 1:50 000 | 1:5000 | | Jackson ImmunoResearch Europe Ltd., Ely, UK/715-005-150 |
| Donkey Anti-Rabbit IgG HRP | 1:50 000 | 1:5000 | | Jackson ImmunoResearch Europe Ltd., Ely, UK/711-035-152 |
| Donkey Anti-Goat IgG HRP | 1:50 000 | 1:5000 | | Jackson ImmunoResearch Europe Ltd., Ely, UK/705-035-003 |
